# Identifiability of Phylogenetic Level-2 Networks under the Jukes-Cantor Model

**DOI:** 10.1101/2025.04.18.649493

**Authors:** Aviva K. Englander, Martin Frohn, Elizabeth Gross, Niels Holtgrefe, Leo van Iersel, Mark Jones, Seth Sullivant

## Abstract

We investigate which evolutionary histories can potentially be reconstructed from sufficiently long DNA sequences by studying the identifiability of phylogenetic networks from sequence data generated under site independent models of molecular evolution. While previous work in the field has established the identifiability of phylogenetic trees and level-1 networks, networks with non-overlapping reticulation cycles, less is known about more complex network structures. In this work, we extend identifiability results to network classes that include pairs of tangled reticulations.

Our main result shows that binary semi-directed level-2 phylogenetic networks are generically identifiable under the Jukes–Cantor model, provided they are triangle-free and strongly tree-child. We also strengthen existing identifiability results for level-1 networks, showing that the number of reticulation nodes is generically identifiable under the Jukes-Cantor model.

In addition, we present more general identifiability results that do not restrict the network level at all and hold for the Jukes-Cantor as well as for the Kimura-2-parameter model. Specifically, we demonstrate that any two binary semi-directed networks that display different sets of 4-leaf subtrees (quartets) are distinguishable. This has direct implications for the identifiability of a network’s reticulated components (blobs). We show that the tree-of-blobs of a network, the global branching structure of the network, is identifiable, as well as the circular ordering of the subnetworks around each blob, for networks in which edges do not cross and taxa are on the outside.

## 1. Introduction

Reticulate events such as hybridization, horizontal gene transfer, and recombination play a central role in shaping the evolutionary histories of many organisms (Arnold, 1997; Barton, 2001; Mallet, 2005, 2007; Keeling and Palmer, 2008; Soucy et al., 2015), including bacteria (e.g. Diop et al., 2022), viruses (e.g. Worobey et al., 2008; Pekar et al., 2021; Jiao et al., 2024), plants (e.g. Rieseberg, 1997; Wang et al., 2024), fish (e.g. Meier et al., 2019), birds (e.g. Taylor and Larson, 2019), and primates (e.g. Patterson et al., 2006; Green et al., 2010; Raghavan et al., 2014). Indeed, due to the importance of such events, phylogenetic networks have emerged as a fundamental framework for modeling evolutionary relationships that involve reticulate events where lineages merge (Nakhleh, 2010; Bapteste et al., 2013; Kong et al., 2022, 2025).

In short, phylogenetic networks are graphs used to represent evolutionary relationships between different taxa, see Figure 1 for an example. A *graph* consists of nodes connected by edges.

**Figure 1.**
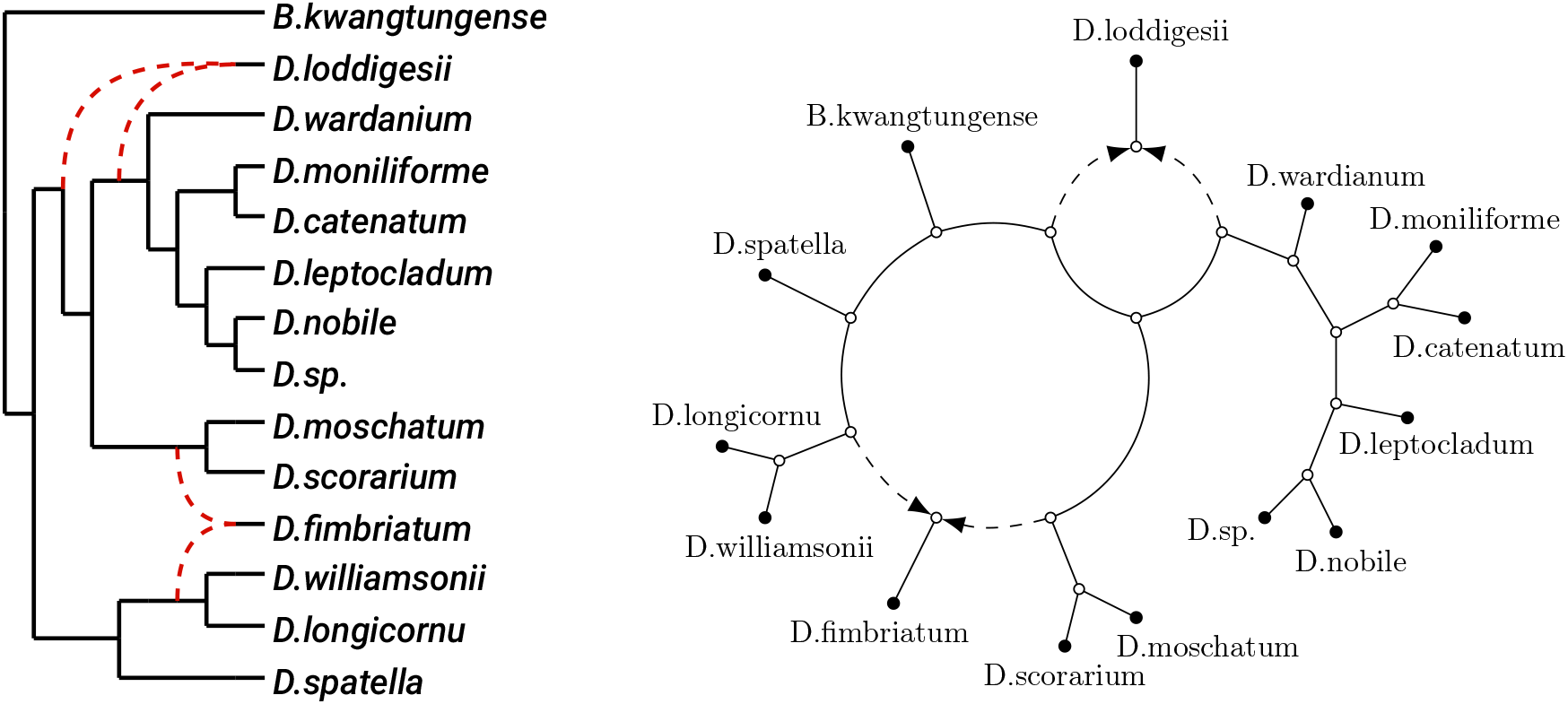
Left: A directed level-2 phylogenetic network of orchids, from Wang et al. (2024), where the directions go from left to right. Right: Its semi-directed network, obtained by suppressing the root and only letting the dashed reticulation edges keep their direction.

The nodes may be observed in the data (the leaves, representing taxa) or inferred by the analysis (internal nodes, representing ancestors). The edges may be directed (from ancestor to descendant) or undirected. Nodes may have more than one parent, making the relationships reticulate. Such nodes are called *reticulation nodes*.

The goal of phylogenetic network inference is to recover these graphs and associated numerical parameters from observed data. Our work assumes that the observed data are a set of aligned DNA sequences, and such sequences evolved from a common ancestor and contain phylogenetic signal that may support reconstruction of the evolutionary history. The objective of our analysis is to determine whether this signal suffices to infer a graph representing an evolutionary history when reticulate events are present.

In fact, as network complexity increases, inferring the generating network may become impossible, even with arbitrarily long sequences. Therefore, a central question is: what network structures are identifiable, i.e., can potentially be uniquely recovered from sufficiently long DNA sequences, assuming a certain model of DNA evolution? This question underlies the development of *consistent* phylogenetic network reconstruction methods. Such methods guarantee recovery of the correct network from perfect data, such as the recently developed methods Nanuq (Allman et al., 2019), TINNIK (Allman et al., 2024a), Nanuq^+^ (Allman et al., 2025b) and Squirrel (Holtgrefe et al., 2025b).

In this paper, we consider identifiability of phylogenetic networks from site-pattern probabilities under a model assuming a Jukes-Cantor (JC) or Kimura-2-Parameter (K2P) substitution process on each network edge, with no incomplete lineage sorting, where, at reticulation nodes, each site is independently inherited from one of the parent nodes under inheritance probabilities. Although this is a heavily simplified model, we see it as a major step towards studying identifiability under more complex models.

To date, most identifiability results and consistent reconstruction methods for phylogenetic networks have been confined to level-1 networks. The breakthrough of this paper is the establishment of identifiability results for level-2 networks, which are significantly more complex than level-1 networks because they allow pairs of “tangled” reticulate events, see Figure 2. This can be made more precise by using the term *blob*, a maximal subnetwork that remains connected after deletion of any single edge. A blob is *trivial* if it consists of a single node. In a level-1 network, each nontrivial blob contains exactly one reticulation node and hence consists of a single cycle. Therefore, any two cycles in a level-1 network must not share an edge. Unfortunately, this restriction excludes many realistic evolutionary scenarios (e.g. Ge et al., 1999; Westenberger et al., 2005; Sessa et al., 2012; Raghavan et al., 2014; Szöllősi et al., 2015; Wang et al., 2024). For example, consider the phylogenetic networks for orchids depicted in Figure 1. The network to the left is directed. Since the root location is not identifiable under many models, we will consider *semi-directed* networks (see the network to the right in Figure 1 and Solís-Lemus and Ané (2016)). Such networks can often be rooted in practice using an outgroup. The networks in Figure 1 are not level-1, since they contain overlapping cycles. They, in fact, belong to a larger class of networks called *level-2* networks, which are binary networks where each blob contains at most two reticulation nodes. Level-2 networks can be seen as networks with isolated reticulations and pairs of tangled reticulations. Tangled reticulations lead to major mathematical and computational challenges, especially when considering identifiability.

**Figure 2.**
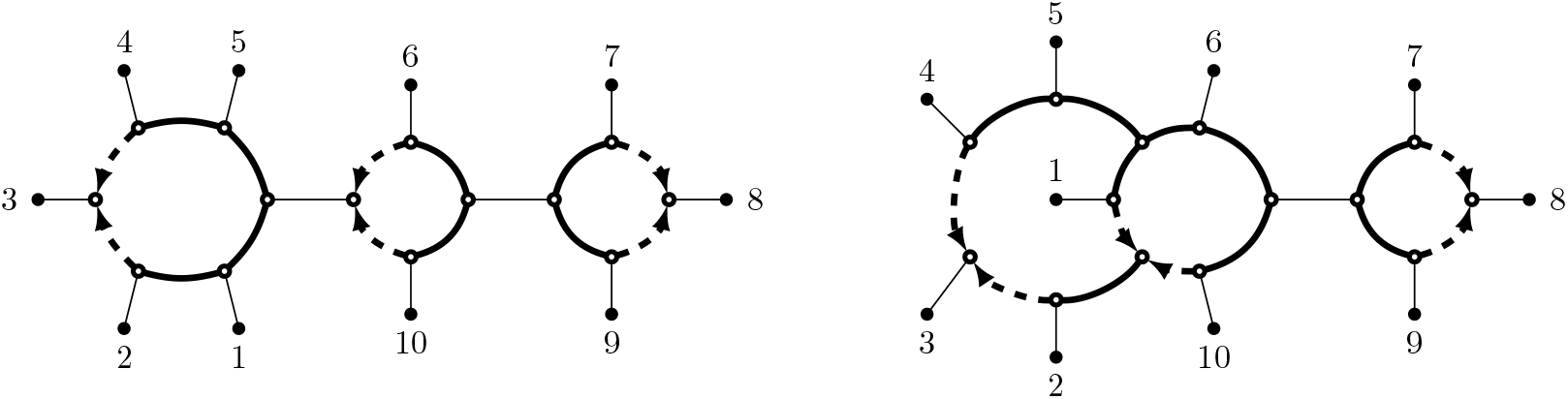
Illustration of the difference between level-1 (left) and level-2 (right) semi-directed phylogenetic networks. Nontrivial blobs are indicated in bold. Dashed arcs are reticulation edges and the nodes they point at are reticulation nodes.

Roughly speaking, our main result is that binary semi-directed level-2 phylogenetic networks are identifiable from site-pattern probabilities under a JC substitution process with inheritance probabilities, assuming generic parameters and under some technical conditions on the network topology, see Theorem 3.2 for a precise statement.

The main advance in Theorem 3.2 is that it allows for tangled pairs of reticulate events. In addition, the result is stated for an arbitrary number of reticulations rather than a fixed number of reticulations (as in Gross et al. (2021) on identifiability of level-1 networks). On the other hand, more complicated structures than level-2 are not allowed, such as triples of reticulations that are all tangled in a single blob. There are two additional technical restrictions on the networks. The first is the triangle-free assumption, which forbids reticulate events between a node and a direct ancestor of that node (leading to a length-3 cycle or *triangle*). Previous results either exclude these scenarios, or in some cases, show that current methods are unable to identify them (Gross and Long, 2018; Baños, 2019; Allman et al., 2022; Xu and Ané, 2023; Allman et al., 2024b). The second is the strongly tree-child condition that forbids a reticulation node to be a direct descendant of another reticulation node (which is known to be unidentifiable (Sullivant, 2025)) and forbids nodes that have two reticulation nodes as direct descendants (less likely scenarios for certain types of data (Kong et al., 2022)).

The proof requires a mix of techniques using tools from algebra and combinatorics. On the algebra side, we find phylogenetic invariants in order to distinguish certain pairs of small networks, and we use a matroid-based approach (Hollering and Sullivant, 2021) to distinguish other pairs. In addition, we derive a new inequality for trinets (3-leaf networks), which can distinguish further pairs of networks under JC. Once we have established a set of distinguishing results for pairs of small networks on 3, 4 and 5 leaves, we are able to use the combinatorial structure of level-2 networks to prove Theorem 3.2. We note that in previous work by Huber et al. (2025), it was shown that all binary semi-directed level-2 networks are uniquely encoded by their induced quarnets (4-leaf networks). However, for the case of binary semi-directed level-2, triangle-free, tree-child networks, it is not true that they can be uniquely recovered from their induced triangle-free, tree-child quarnets, as we will later see in Figure 6. Thus, this requires us to prove some distinguishing results beyond just quarnets and beyond just triangle-free tree-child quarnets.

As we build to our main theorem, we prove several other results that are interesting in their own right and hold for networks of any level. First of all, we show that under JC and K2P, if two networks have different sets of displayed quartet trees, then they are distinguishable. This has consequences for the identifiability of the *tree-of-blobs*, the tree obtained from a network by contracting each blob to a node (Gusfield et al., 2007; Allman et al., 2023, 2024a), which captures the global branching structure of a phylogenetic network, see Figure 3. It is important to note that reconstructing a tree-of-blobs is fundamentally different from reconstructing a phylogenetic tree. Although the tree-of-blobs is a tree, it is based on evolutionary models that allow reticulation rather than assuming a purely treelike history. By combining our result with results from Allman et al. (2023); Huber et al. (2025); Rhodes et al. (2025), it follows that, under JC and K2P, the tree-of-blobs of any binary semi-directed network is identifiable as well as the circular ordering of the subnetworks around certain blobs (see Theorem 3.3).

**Figure 3.**
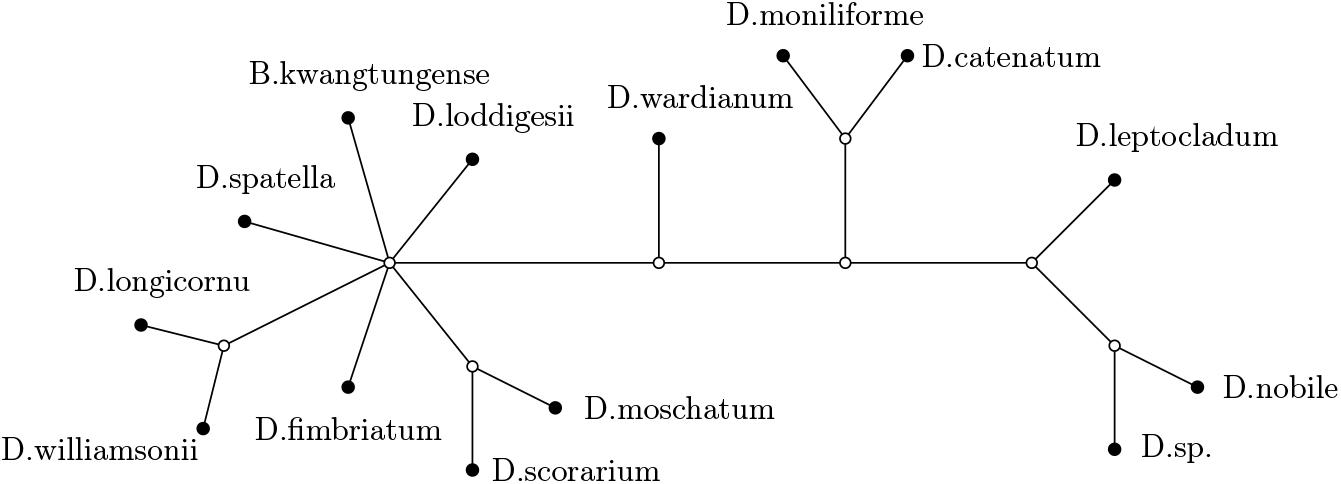
The tree-of-blobs of the semi-directed level-2 network on 14 orchids from Figure 1.

We also improve the results of Gross et al. (2021), by strengthening the identifiability results for binary level-1 networks under JC. We show that we do not need to assume that the networks are triangle-free, as long as we do not need to identify the reticulation node in a triangle. Moreover, we show that one can even distinguish between level-1 networks with different numbers of reticulate events (see Theorem 3.1). This could be important since the number of reticulate events is in practice usually not known in advance. In particular, it means that we can distinguish between a tree and a non-tree level-1 network. We show that this also holds for level-2 networks and conjecture that trees can be distinguished from non-trees for all levels (see Conjecture 2.16 for the precise technical version).

### 1.1. Related work on the identifiability of phylogenetic networks

A prominent line of research focuses on network inference from gene tree distributions, where gene trees arise under the multispecies coalescent process (Yu et al., 2011). In this context, the problem of identifiability is analyzed using gene tree probabilities and related summary statistics such as quartet concordance factors. This line of research (Solís-Lemus and Ané, 2016; Baños, 2019; Allman et al., 2023; Rhodes et al., 2025; Allman et al., 2024b) eventually led to a proof of the identifiability of semi-directed level-1 phylogenetic networks up to the orientation of triangles, the identifiability of the tree-of-blobs of general networks and of the circular ordering around certain blobs, from quartet concordance factors. Very recently, Allman et al. (2025a) proved identifiability for the class of non-binary, strongly tree-child networks that are *galled* (reticulations have no ancestral reticulations within the same blob), subject to further topological restrictions related to cycle sizes. Moreover, Holtgrefe et al. (2025a) obtained identifiability for galled level-2 networks under a planarity condition called *outer-labeled planar* (see Definition 2.3), without the strongly tree-child condition but only up to a canonical form.

On the other hand, there have been a number of papers which specifically consider Markov models of evolution of aligned sequence data on a network. So far, such work on networks usually assumes that each column of the alignment evolves independently down a tree displayed by the network, contrasting with coalescent-based models which consider all trees. Under these models, parallel edges are not identifiable (Gross and Long, 2018), nor are blobs with only two incident edges (Sullivant, 2025). Biologically, this corresponds to scenarios in which reticulated activity eventually produces just one lineage, which does not provide enough information to reconstruct the reticulate events under these models. Since such scenarios are not identifiable, we will exclude them throughout this paper.

Gross and Long (2018) proved, using algebraic geometry, that binary large-cycle networks, which are binary semi-directed phylogenetic networks with a single cycle of length greater than three, are generically identifiable under JC. Hollering and Sullivant (2021) used algebraic matroids to show that pairs of phylogenetic trees are generically identifiable for 2-tree CFN and Kimura 3-Parameter (K3P) mixtures, where each column evolves down one of two trees, and that binary large-cycle networks are generically identifiable under K2P and K3P. Gross et al. (2021) showed that binary semi-directed level-1 networks are generically identifiable under JC, K2P and K3P, assuming the network is triangle-free and the number of reticulation nodes is known. While these works mostly focused on level-1 networks, Ardiyansyah (2021) explored an acutely restricted class of level-2 networks in a similar algebraic setting.

Furthermore, Francis and Moulton (2018) considered a different extension of the tree-based Markov model to networks, where blocks of columns of the alignment evolve down the same displayed tree and there is a fixed probability to start a new block evolving down another displayed tree. For this model they showed an identifiability result for the class of rooted binary tree-child networks with the same number of reticulation nodes, at the cost of fixing the parameter values of the evolutionary model.

Allman et al. (2022) use yet a different approach by showing that semi-directed level-1 networks are identifiable from sequence data via inter-taxon logDet distances under a model that includes incomplete lineage sorting and with some further restrictions on the networks.

Another very recent research direction considers the number of sites needed to correctly reconstruct a phylogenetic network with high probability (Frohn et al., 2026), which has so far also been restricted to level-1 networks.

### 1.2. Outline of the paper

In the next section, we cover preliminary background on networks, Markov processes on trees and networks, and identifiability including the algebraic approaches to identifiability we will use. In Section 2.2, we use linear invariants and inequalities to show that under JC and K2P, two networks that have different displayed quartet trees are distinguishable. In Section 2.3, we use phylogenetic invariants and matroids to distinguish certain pairs of quarnets (4-leaf subnetworks) which will be used in later proofs. In Section 2.4, we derive a new trinet inequality that can distinguish a 3-leaf tree from a level-1 or level-2 trinet (3-leaf subnetwork under JC. Then, in the Results section, we put all these results together to prove our main results described above. Appendix A contains the proofs omitted from Sections 2.2, 2.3, 2.4 and 3.2 due to their length. We end with a Discussion section.

## 2. Materials & Methods

### 2.1. Preliminaries

#### 2.1.1. Phylogenetic networks

In this section, we introduce concepts from graph theory that are utilized to study phylogenetic networks. The *lowest stable ancestor* in a rooted directed acyclic graph is the lowest node through which all directed paths from the root to a leaf pass. The *in-degree* of a node *v* is the number of edges that is incident to *v* and directed towards *v* while the *out-degree* of *v* is the number of incident edges directed away from *v*.

##### Definition 2.1

A *directed (binary phylogenetic) network N* ^+^ on a set of taxa X is a directed acyclic graph where

i. X is the set of nodes with out-degree zero and in-degree one (*leaves*);
ii. there is a unique node with in-degree zero (the *root*), which has out-degree two and is the lowest stable ancestor; and
iii. all other nodes have in-degree one and out-degree two (*tree nodes*) or in-degree two and out-degree one (*reticulation nodes*).

Note that such a network is connected since it is acyclic and has a unique root. Parallel edges are not allowed. The network *N* ^+^ is a *rooted (binary) phylogenetic tree* if the network does not contain any reticulation nodes. We call a non-leaf node in *N* ^+^ *internal* and an edge directed into a reticulation node a *reticulation edge*. A pair of reticulation nodes *v, w* is called *stacked*, if one is a child of the other.

##### Definition 2.2

A *semi-directed (binary phylogenetic) network N* on a set of taxa *X* is a partially directed graph that can be obtained from a directed binary phylogenetic network on *X* by removing edge directions of non-reticulation edges and suppressing the former root.

Reticulation nodes, reticulation edges, internal nodes, and leaves are defined analogously for semi-directed networks. Also in these networks we do not allow parallel edges. A semi-directed network without reticulation nodes is called an *unrooted (binary) phylogenetic tree*. Such a tree is *displayed* by a semi-directed network on *X*, if it can be obtained by deleting exactly one incoming reticulation edge per reticulation node, omitting the direction of the other incoming edge, and then exhaustively deleting out-degree-0 nodes not in **X** and suppressing degree-2 nodes. See Figure 4 for an example of all unrooted phylogenetic trees displayed by a semi-directed network.

A node *v* is *below* a reticulation node *r* in a semi-directed network *N* if there is a path from *r* to *v* in which all traversed reticulation edges are directed towards *v*. The node *v* is *directly below r* if *v* and *r* are also adjacent. Every *cut-edge e* of a (semi-)directed network, i.e. an edge whose removal disconnects the network, induces a bi-partition *A* ∪ *B* of its leaves *X* with *A, B*≠ ∅, *A* ∩ *B* = ∅, in which case we say that *e* induces the *split A*|*B*. The split is *nontrivial* if both |*A*|, |*B*| ≥ 2.

A *blob* of a (semi-)directed phylogenetic network is a maximal connected subgraph without any cut-edges. Such a blob is an *m*-blob for some positive integer *m* if the blob is incident to exactly *m* edges not in the blob. A blob is *trivial* if it consists of a single node. The unrooted (possibly non-binary) phylogenetic tree that is obtained from a semi-directed network by contracting every blob to a single node, and suppressing degree-2 nodes, is called the *tree-of-blobs*.

**Figure 4.**
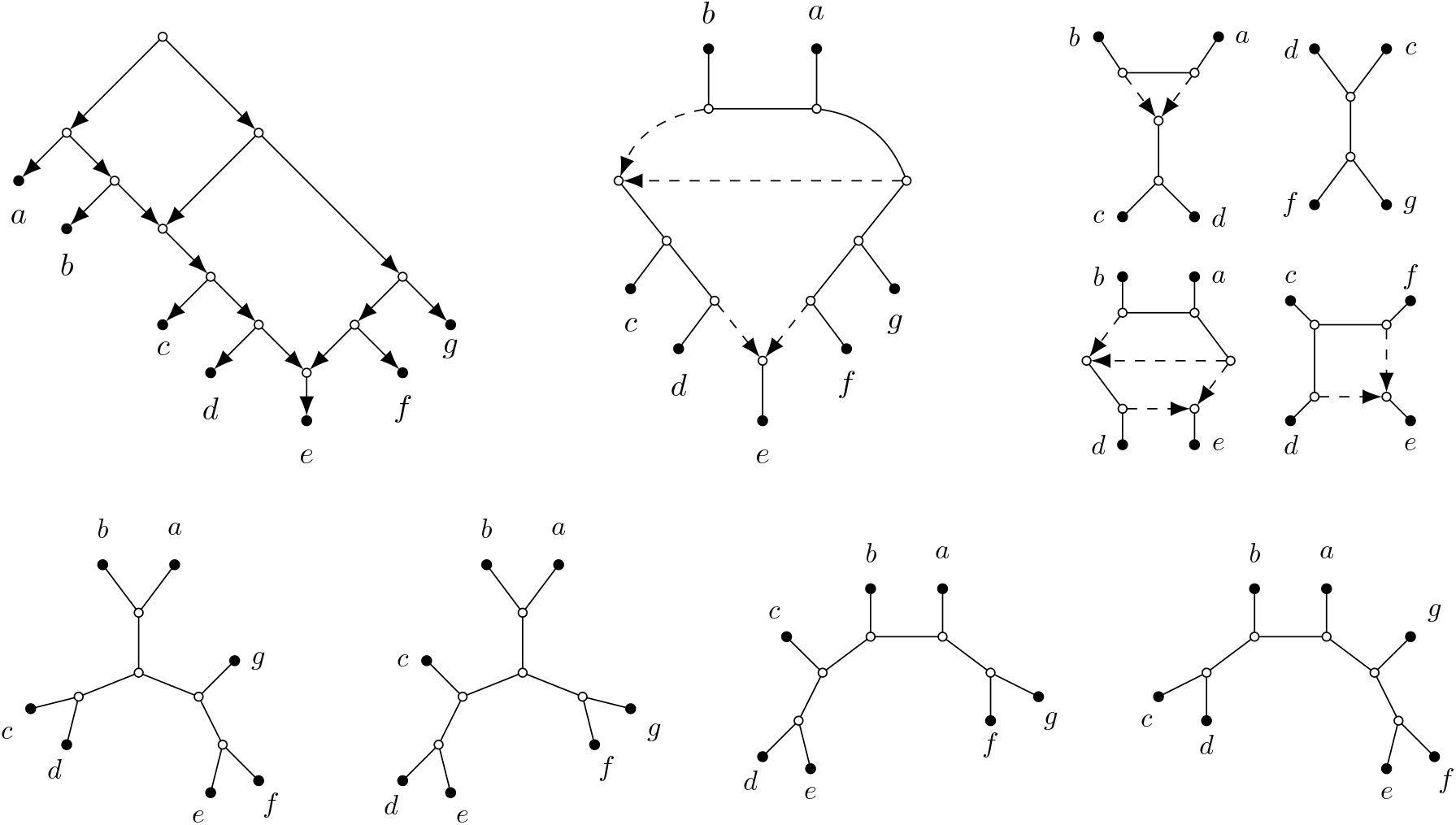
*Top left:* A directed phylogenetic network *N* ^+^ on *X* = {*a, b, c, d, e, f, g*}. *Top middle:* The semi-directed phylogenetic network *N* obtained from *N* ^+^. *Top right:* Four of the quarnets induced by *N*. *Bottom:* The four unrooted phylogenetic trees on *X* displayed by *N*.

##### Definition 2.3

Let *N* ^+^ be a directed phylogenetic network (respectively, *N* a semi-directed phylogenetic network), and let *k* be some non-negative integer. Then, we call the network *N* ^+^ (respectively, *N*)

i. *level-k* if there are at most *k* reticulation nodes in each blob of the network;
ii. *strict level-k* if the network is level-*k* but not level-(*k* − 1);
iii. *stack-free* if none of the reticulation nodes of the network are stacked;
iv. *simple* if the network has a single non-leaf blob;
v. *triangle-free* if the network contains no cycles of length three, where edge directions are disregarded;
vi. *outer-labeled planar* if the network has a planar embedding without crossing edges and with the leaves on the outside.

Furthermore, *N* ^+^ is *tree-child* if every internal node has at least one child that is either a tree node or a leaf. *N* is *strongly tree-child* if every directed network from which *N* can be obtained (as in Definition 2.2) is tree-child. Holtgrefe et al. (2026) showed that *N* is strongly tree-child if and only if each node with an outgoing edge also has two incident undirected edges.

To illustrate some of these definitions, consider Figure 4, which depicts a tree-child directed network *N* ^+^ and the strongly tree-child semi-directed network *N* that can be obtained from *N* ^+^. Note that, while the directed network *N* ^+^ is not the unique directed phylogenetic network from which *N* can be obtained, all possible rootings of *N* result in a tree-child network. Both networks are strict level-2, simple, triangle-free, stack-free and outer-labeled planar. By contrast, the semi-directed network in Figure 1 is not simple because every internal node not included in a cycle is also a blob, i.e., the network contains eight non-leaf blobs. Also note that our definition of a simple network is different from the definition of simple graphs in standard graph theory terminology. The semi-directed network on the right in Figure 2 is an example of a network that is not outer-labeled planar.

*Remark*. Throughout this work we assume that semi-directed networks have no 2-blobs or parallel edges, since they can be safely suppressed under our considered models, see Sullivant (2025); Gross and Long (2018). Note that the absence of such substructures is implied by the tree-child condition. Moreover, all our identifiability results extend to the setting where such structures are allowed but one aims to identify the network with these structures suppressed. Note that this property is specific to *equivariant models* (which include the JC and K2P models we consider) and does not extend to more general models such as the GTR model.

Since outer-labeled planar, semi-directed networks can be drawn in the plane with leaves on the outside, traversing the boundary of such a drawing yields a *circular order* (i.e., an order up to reversal and cyclic permutations) of the leaves. In general, an outer-labeled planar network may have multiple such drawings, and consequently multiple such *congruent* circular orders. However, as shown by Rhodes et al. (2025), if the network is also simple, this congruent circular order is unique. For example, the network *N* in Figure 4 is congruent with only the circular order (*a, b, c, d, e, f, g*).

An *up-down path* between two leaves *x*_*i*_ and *x*_*j*_ of a semi-directed network is a path of *k* edges where the first *l* edges are directed towards *x*_*i*_ and the last *k* − *l* edges are directed towards *x*_*j*_, for some 0 ≤ *l* ≤ *k*, considering undirected edges to be bidirected. Using this notion, we can formally define *induced subnetworks* of a semi-directed network.

##### Definition 2.4

Given a semi-directed network *N* on *X* and *X* ⊂ *X* with |*X*| ≥ 2, we define the *subnetwork* of *N induced* by *X* as the semi-directed network *N* _*X*_ obtained from *N* by taking the union of all up-down paths between leaves in *X*, followed by exhaustively suppressing all nodes of degree two, parallel edges and 2-blobs.

If |*X* | = 3, |*X*| = 4 or |*X*| = 5, then we call *N* | _*X*_ a *trinet, quarnet* or *quinnet* induced by *N*, respectively. See Figure 4 for an example of four quarnets induced by a semi-directed network. Proving identifiability by restricting to (level-2) trinets, quarnets, and quinnets will be a key idea in our proofs. Figure 5 shows all sixteen semi-directed simple level-2 quarnets, up to labeling the leaves. Whenever we refer to a quarnet of a specific type, we use the annotations as in this figure. For example, the quarnet of type 0 is the only simple quarnet that is strict level-1.

**Figure 5.**
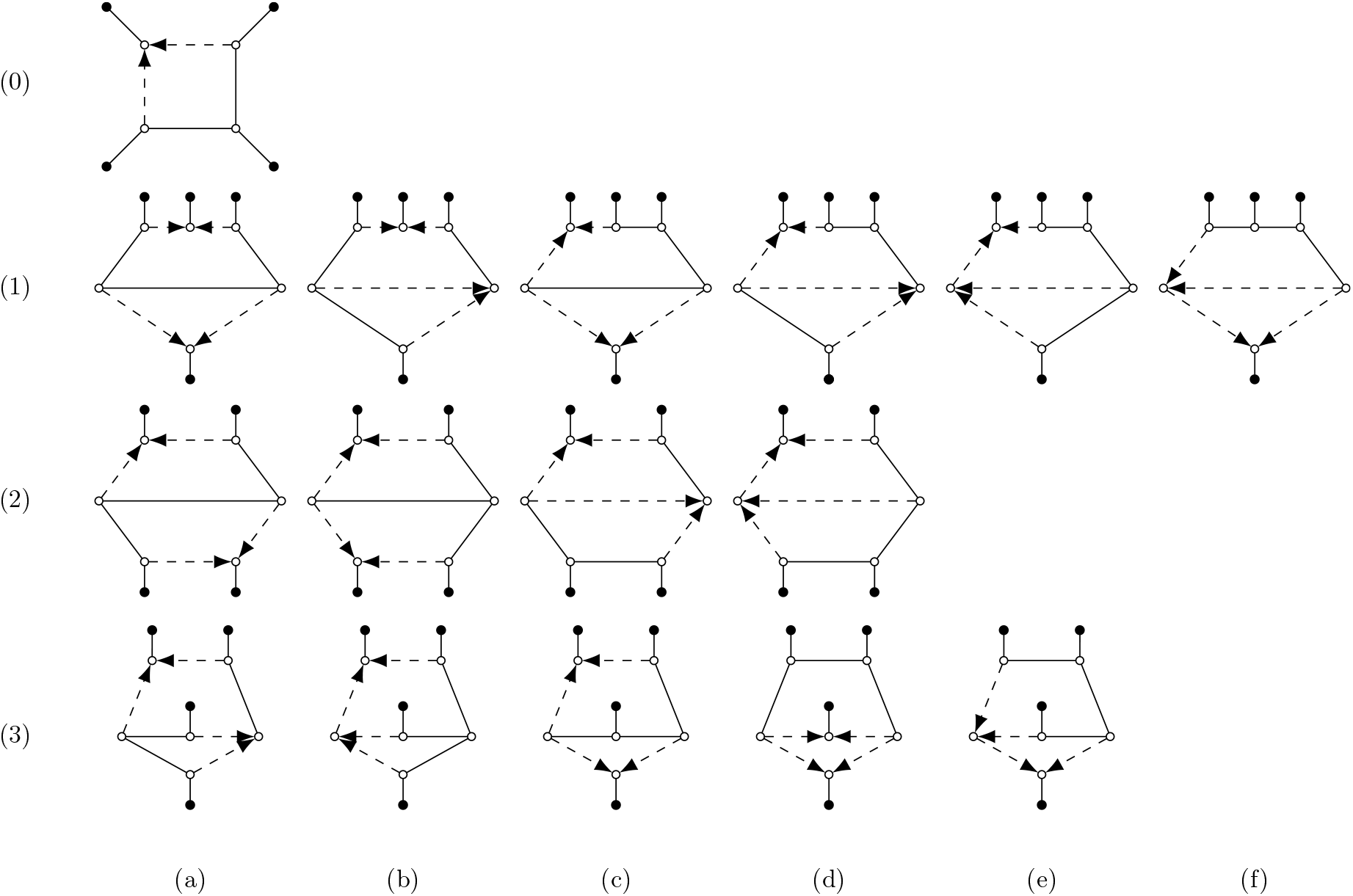
All sixteen semi-directed simple level-2 quarnets up to labeling the leaves.

#### 2.1.2. Network-based Markov models

In this paper, we consider network-based Markov models, which are described most naturally via a directed phylogenetic network. However, for reversible models, such as the JC and K2P models considered here, any two directed networks 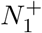 and 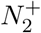 that have the same semi-directed network *N* will yield the same set of probability distributions (see proof of Proposition 1 in Gross et al. (2021)). Thus, we can use directed networks and semi-directed networks interchangeably when describing model parametrizations. We will give the definitions for directed networks since this is more convenient.

Given an *n*-leaf directed phylogenetic network *N* ^+^ = (*V, E*),we associate with each node *v* ∈ *V* a random variable *X*_*v*_ with state space Σ = *{A, G, C, T}*, corresponding to the four DNA bases.

The JC and K2P substitution models then assume a uniform distribution *π* at the root, and for each edge *e* = (*u, v*), a transition matrix *M* ^*e*^ governing the probabilities of nucleotide change from *u* to *v* (Jukes and Cantor, 1969; Kimura, 1980). This yields joint probability distributions *p*_*ω*_(*T*) of a site pattern *ω* ∈ Σ^*n*^ observed at the leaves in **X** of each *switching T* of *N* ^+^. Here, a switching of *N* ^+^ is a tree (possibly having degree-2 nodes and additional leaves not in *X*) obtained by deleting exactly one incoming reticulation edge per reticulation node. See, for example, Gross et al. (2021) for more details.

To derive the site pattern probability distribution of *N*^+^, let *k* be the number of reticulation nodes of *N* ^+^ and consider the set 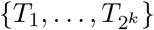 of switchings of *N* ^+^. Assign a parameter δ_*i*_ ∈ (0, 1) to each reticulation node *r*_*i*_ with corresponding reticulation edges 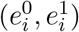. Here, δ_*i*_ (resp. 1 − δ_*i*_) is an *inheritance probability*, representing the probability that a particular site was inherited along edge 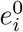 (resp.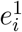). Note that we do not allow inheritance probabilities equal to 0 or 1. For *j* ∈ *{*1, …, 2^*k*^*}*, let ζ_*j*_ ∈ *{*0, 1*}*^*k*^ denote the indicator variable for the reticulation edges 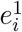 present in *T*_*j*_. Then, the probability of observing site pattern *ω* in *N* ^+^ is given by

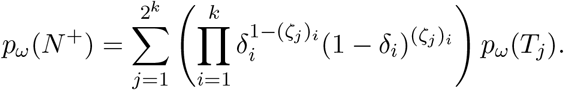

Thus, for the stochastic parameter space 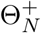 of transition matrices *M* ^*e*^, inheritance probabilities δ_*i*_ and root distribution *π*, we can view both the JC and K2P model on *N* ^+^ as the image of the polynomial map

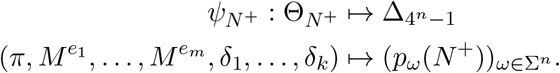

The image of map 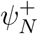 is denoted 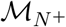 and called the *phylogenetic network model (associated to N* ^+^) or *JC network model* or *K2P network model* if, respectively, JC or K2P constraints on the substitution processes are used.

For the JC and K2P models, a linear change of coordinates — the Fourier (or Hadamard) transform (Hendy, 1989; Hendy and Penny, 1989; Evans and Speed, 1993) — simplifies the parameterizations. We describe the parameterization of this model in Fourier coordinates *q*_*ω*_(*T*) following Sturmfels and Sullivant (2005). First, we identify our state space Σ with the Klein four-group z_2_ *×* z_2_ as follows: *A* = (0, 0), *G* = (1, 0), *C* = (0, 1), *T* = (1, 1). We associate four Fourier parameters with each edge *e* ∈ *E*, denoted by 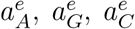 and 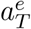. In addition, under the JC model we have that 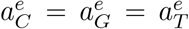, and under the K2P model we have that 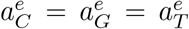. Furthermore, 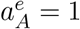 for both models. We assume throughout this paper that all other parameters are in the interval (0, 1), which corresponds to excluding zero and infinite branch lengths. Then, for site pattern *ω* = (*g*_1_, *g*_2_, …, *g*_*n*_), the *Fourier coordinates* of *p*_*ω*_(*T*) are given by

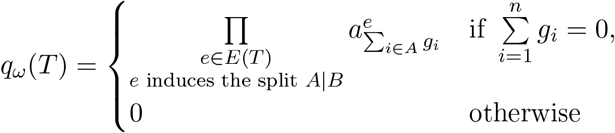

where addition is in the group z_2_ *×* z_2_. For any edge *e* ∈ *E* which induces the split *A*|*B*, we also write Fourier parameter 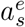 as 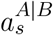 for all *s* ∈ Σ.

We summarize that for the JC and K2P models, the transformed probability distribution for each site-pattern in a phylogenetic tree model can be parameterized by a monomial with one parameter per edge in the tree. Therefore, transformed distributions from a phylogenetic network model can be parameterized by a sum of monomials, one for each tree *T*_*j*_, *j* ∈ {1, …, 2^*k*^}, weighted by the corresponding inheritance probabilities.

#### 2.1.3. Generic identifiability and distinguishability

Our paper seeks to answer questions about the distinguishability and generic identifiability of phylogenetic networks, which we define and discuss here.

##### Definition 2.5

Let *{M*_*N*_ *}*_*N*∈*N*_ be a finite class of phylogenetic network models and 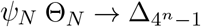 be the parameterization map for a phylogenetic network model. Given two distinct networks *N*_1_ and *N*_2_ on the same set of *n* leaves, we say they are *distinguishable* if the set of numerical parameters in 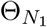 that 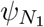 maps into 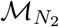 and the set of numerical parameters in 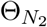 that 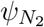 maps into 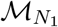 both have Lebesgue measure zero. Then, the network parameter of a phylogenetic network model is *generically identifiable* (w.r.t. the class *N*) if given any two distinct *n*-leaf networks *N*_1_, *N*_2_ ∈ *N*, the networks *N*_1_ and *N*_2_ are distinguishable.

One of our main tools for proving identifiability results for a phylogenetic network model *M*_*N*_ will be to look at its *vanishing ideal* or *ideal of phylogenetic invariants*, defined as the set of polynomials

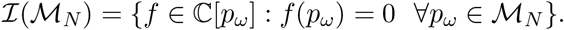

Here, we consider the polynomial ring C[*p*_*ω*_] over indeterminates *p*_*ω*_, with *ω* ∈ Σ^*n*^. Clearly, each phylogenetic invariant *f* vanishes on all probability distributions of a given phylogenetic model.

In particular, we have the following result that relates the existence of phylogenetic invariants to distinguishability (Sullivant, 2023).

##### Proposition 2.6

*Let* 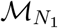 *and* 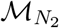 *be two phylogenetic network models on the same number of leaves. Suppose that* 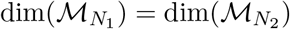. *If there exist a polynomial* 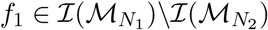 *then networks N*_1_ *and N*_2_ *are distinguishable*.

It is useful to note that when 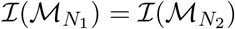, this is not enough to prove that models 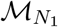 and 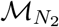 are not distinguishable. In these cases, further methods, such as using semialgebraic constraints, are necessary to prove distinguishability. The following works use semialgebraic methods for identifiability for the interested reader: Allman et al. (2014); Kosta and Kubjas (2017); Casanellas et al. (2021); Allman et al. (2024b).

Since computing phylogenetic invariants can be challenging, the following alternative method based on algebraic matroids — derived from Proposition 3.1 and Algorithm 3.3 in Hollering and Sullivant (2021) — can also be used to establish network distinguishability.

##### Proposition 2.7

*Let* 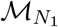 *and* 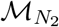 *be two phylogenetic network models on the same number of leaves, and suppose that* 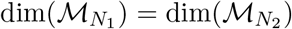. *For i* ∈ *{*1, 2*}, let* 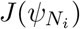 *be the Jacobian matrix of* 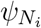 *whose rows and columns are indexed by site patterns* Σ *and model parameters, respectively. Suppose there is a subset S* ⊆ Σ^*n*^ *such that*

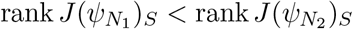

*where J*(*ψ*_*N*_)_*S*_ *denotes the submatrix of J*(*ψ*_*N*_) *consisting of the rows indexed by S. Then, networks N*_1_ *and N*_2_ *are distinguishable*.

Additionally, the following proposition shows that to prove distinguishability, it suffices to consider subnetworks induced by subsets of leaves. See Proposition 4.3 of Gross and Long (2018) for a related result.

##### Proposition 2.8

*Let N*_1_ *and N*_2_ *be two semi-directed networks on the same leaf set X and S* ⊂ *X. If N*_1_|_*S*_ *and N*_2_|_*S*_ *are distinguishable, then N*_1_ *and N*_2_ *are distinguishable*.

*Proof*. Since our models are expressed as the images of polynomial maps, the set of parameter values that map 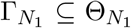 into 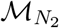 will be a semialgebraic set. That set will have measure zero inside 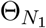 if and only if it is a semialgebraic set of strictly smaller dimension than 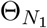. So taking the image of 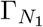 by polynomial map 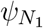 must also give a subset of 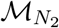 of strictly smaller dimension. Hence, for algebraic phylogenetic models, we have the following equivalent definition of distinguishability: networks *N*_1_ and *N*_2_ are distinguishable if 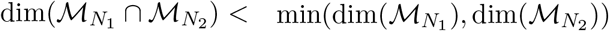. This equivalence will allow us to use previous results directly.

For *i* ∈ {1, 2}, let *N*_*i*_(*S*) be obtained by taking the union of all up-down paths in *N*_*i*_ between pairs of leaves in *S* (keeping degree-2 nodes and 2-blobs). If *N*_1_ | _*S*_, *N*_2_ | _*S*_ are distinguishable, then *N*_1_ | _*S*_, *N*_2_(*S*) are distinguishable, so *N*_1_(*S*), *N*_2_(*S*) are distinguishable (Gross and Long, 2018; Sullivant, 2025).

We now show that it follows that *N*_1_ and *N*_2_ are distinguishable. Consider the marginalization over the states of **X* \ S* given by a linear map 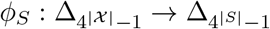, where 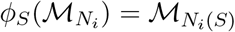. In addition, we have a coordinate projection map 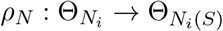 which keeps the parameters that appear in *N*_*i*_(*S*). Then, when we consider 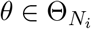 such that 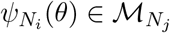, we get

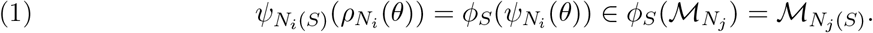

Since *N*_1_(*S*) and *N*_2_(*S*) are distinguishable, for *i, j* ∈ *{*1, 2*}, i ≠ j*, the set of numerical parameters 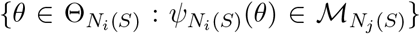 is a set of Lebesgue measure zero. Since 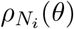 projects a finite number of parameters out, we conclude that the set of numerical parameters in 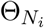 that 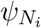 maps into 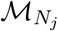 has Lebesgue measure zero, because the preimage under a surjective polynomial map of a set of measure zero also has measure zero. Thus, *N*_1_ and *N*_2_ are distinguishable.

With Proposition 2.8 in mind, our strategy for proving identifiability of the network parameter for our considered class of networks becomes clear. We will show distinguishability of many pairs of small networks, using different strategies including computing phylogenetic invariants, matroid calculations, and finding inequalities that can distinguish the small networks. Then we will show that arbitrary size networks can be distinguished by considering subnetworks induced by small sets of leaves. In particular, in the next section we will provide semialgebraic sufficient conditions under which specific phylogenetic network models do not intersect which we will make use of repeatedly to prove distinguishability throughout the rest of the paper.

### 2.2. Distinguishing Networks with Different Displayed Quartets

This section builds on the linear invariants and inequalities developed by Allman et al. (2010, 2012) for tree mixture models to address identifiability of networks under the JC and K2P models. We show that networks of any level with different displayed quartets are distinguishable, and as a corollary, the tree-of-blobs of a network is also identifiable.

#### Proposition 2.9

*Consider the Fourier coordinates q associated to a quartet tree T with leaf set {*1, 2, 3, 4*} under the JC or K2P model. Then, the invariant*

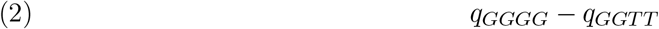

*evaluates to zero if T has split* 12|34 *and is strictly positive if T has split* 13|24 *or* 14|23. *Proof*. In the first case, we have that

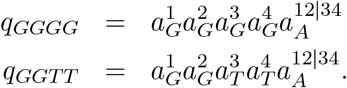

However, since 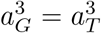 and 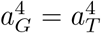 for JC and K2P we see that expression 2 evaluates to 0. On the other hand, for the tree with split 13|24 we have

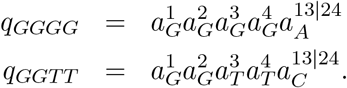

Then, since 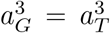 and 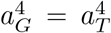, and 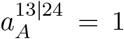 for JC and K2P, expression 2 becomes 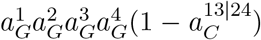. Since 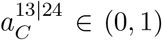, this is positive. A similar calculation shows that expression 2 is positive for distributions that come from the tree with split 14|23.

The proof of the following result is along similar lines to the proof of Proposition 2.9 and is thus deferred to the appendix.

#### Proposition 2.10

*Consider the Fourier coordinates q associated to a quartet tree T with leaf set {*1, 2, 3, 4*} under the JC or K2P model. Then, the invariant*

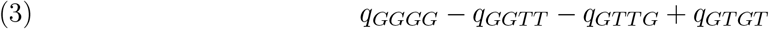

*evaluates to zero if T has split* 12|34 *or* 14|23 *and is strictly positive if T has split* 13|24.

#### Theorem 2.11

*Let N*_1_ *and N*_2_ *be two binary semi-directed phylogenetic networks on the same leaf set *X*. Let* 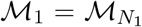 *and* 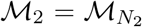 *be the associated JC or K2P network models*. *If there is a* 4 *element subset S* ⊆ *X such that the quarnets N*_1_|_*S*_ *and N*_2_|_*S*_ *have different sets of displayed quartets, then M*_1_ ∩ *M*_2_ = ∅.

*M* ∩ *M* ∅ *M*

*Proof*. Due to Proposition 2.8 it suffices to assume that *N*_1_ and *N*_2_ are themselves quarnets, since if the marginalizations are disjoint then *M*_1_ ∩ *M*_2_ = ∅. Let *d*_*i*_ be a distribution in *M*_*i*_ and note that it is a mixture of distributions from the switchings, or equivalently, from the displayed trees. Let *D*_*i*_ be the set of displayed quartets from network *N*_*i*_. We will assume that *D*_1_ = *D*_2_, and show that *d*_1_ = *d*_2_, which proves *M*_1_ ∩ *M*_2_ = ∅ if all mixing weights are positive.

Suppose that one of *D*_1_ or *D*_2_ is a singleton. After symmetry, we can assume that *D*_1_ = {12 | 34} and that *D*_2_ contains one of the other quartet trees. By Proposition 2.9, expression 2 evaluates to 0 for *d*_1_ and is positive for *d*_2_, so *d*_1_ /= *d*_2_.

Now assume that *D*_1_ has two quartet trees, and *D*_2_ has two or three trees. After symmetry, we can assume that *D*_1_ = *{*12|34, 14|23*}* and that *{*13|24*}* ⊆ *D*_2_. Then, by Proposition 2.10, expression 3 evaluates to 0 for *d*_1_, whereas it is positive for *d*_2_. So, again, *d*_1_≠ *d*_2_.

In the following corollary we show that the tree-of-blobs of a semi-directed network of any level is identifiable under the JC and K2P network models.

#### Corollary 2.12

*Let N*_1_ *and N*_2_ *be two binary semi-directed phylogenetic networks on the same leaf set X. Let* 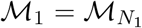 *and* 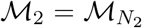 *be the associated JC or K2P network models. If the trees-of-blobs of N*_1_ *and N*_2_ *are not isomorphic, then M*_1_ ∩ *M*_2_ = ∅.

*Proof*. In Huber et al. (2025) it is shown that the tree-of-blobs of a semi-directed network of any level is uniquely determined by the nontrivial splits of its quarnets. Since *N*_1_ and *N*_2_ have non-isomorphic trees-of-blobs, for some 4 element set *S* ⊆ *X* the quarnets *N*_1_ | _*S*_ and *N*_2_ | _*S*_ do not induce the same nontrivial split (with possibly one of them not inducing a nontrivial split at all). Then, by Proposition A.1a, the two quarnets display a different quartet. The result then follows from Theorem 2.11.

#### Corollary 2.13

*Let N*_1_ *and N*_2_ *be two binary semi-directed phylogenetic networks on the same leaf set X. Let* 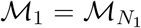 *and* 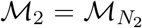 *be the associated JC or K2P network models. If N*_1_ *and N*_2_ *are both outer-labeled planar and they have different sets of congruent circular orders, then M*_1_ ∩ *M*_2_ = ∅.

*Proof*. By the previous corollary, we may assume without loss of generality that *N*_1_ and *N*_2_ have the same tree-of-blobs. Then, for the two networks to have different sets of congruent circular orders, there must be an *m*-blob with *m* ≥ 4 in the two networks which induces a different ordering of the subnetworks around it in *N*_1_ than in *N*_2_. Hence, we may assume without loss of generality that *N*_1_ and *N*_2_ are both simple networks, i.e. they consist of a single non-leaf blob. In Theorem 5.3 of Rhodes et al. (2025) it is shown that the circular order of a binary, simple, outer-labeled planar semi-directed phylogenetic network is uniquely determined by the circular orders of all its simple, outer-labeled planar quarnets. Since we assume that *N*_1_ and *N*_2_ are congruent with a different circular order, there is some 4-leaf set *A* ⊆ **X** such that *N*_1_ | _*A*_ is congruent with a unique circular order and *N*_2_ | _*A*_ is congruent with a different unique circular order or it is not congruent with a unique circular order. Then, by Proposition A.1, the two quarnets display different quartets. The result then follows from Theorem 2.11.

### 2.3. Distinguishing Small Networks using Invariants and Matroids

In this section, we describe a lemma that distinguishes pairs of quarnets, which will later support identifiability of larger networks. The results in this section are computational: Lemma 2.14 is verified by a sequence of `Macaulay2` calculations (Grayson and Stillman, 1992). Direct calculations of the vanishing ideals of our considered models seem out of reach at present, so the computations instead rely on the underlying ideas of Propositions 2.6 and 2.7.

#### Lemma 2.14

*Let N*_1_ *and N*_2_ *be two binary level-2 semi-directed phylogenetic networks on the same leaf set *X*. Let S be a 4 leaf subset of *X*. Then, under the JC model:*

a. *if N*_1_ | _*S*_ *and N*_2_ | _*S*_ *are distinct quarnets of type 1b with a different leaf in the top middle in the planar representation of Figure 5, then N*_1_ *and N*_2_ *are distinguishable;*
b. *if N*_1_ | _*S*_ *and N*_2_ | _*S*_ *are distinct quarnets of type 2a in Figure 5, then N*_1_ *and N*_2_ *are distinguishable;*
c. *if N*_1_ | _*S*_ *and N*_2_ | _*S*_ *are distinct quarnets of type 3a, 3c or 3d in Figure 5, then N*_1_ *and N*_2_ *are distinguishable*. *Let S* = *{a, b, c, d, e} be a 5 leaf subset of X. Then, under the JC model:*
d. *if N*_1_|_*S*_ *and N*_2_|_*S*_ *are the two quinnets from Figure 6, then N*_1_ *and N*_2_ *are distinguishable*.

*Proof*. For each of these results, we may omit the |_*S*_ notation, since it suffices to consider the case that *S* is the set of leaves of the networks under consideration, simply by marginalizing (see Proposition 2.8).

The computational proofs of these results were carried out in `Macaulay2` (Grayson and Stillman, 1992), using the algorithm available through (Grindstaff et al., 2025) to obtain the parameterizations. Parts (a), (b) and (d) employ the matroid method of comparing the ranks of submatrices of the Jacobians from the two networks (see Proposition 2.7). For part (c) we instead compute low degree invariants of our networks using the *MultigradedImplicitization* toolbox (Cummings and Hollering, 2025) in `Macaulay2`, with distinguishability following from Proposition 2.6. In Appendix A, we discuss the files and explain how each of these computations work in detail.

**Figure 6.**
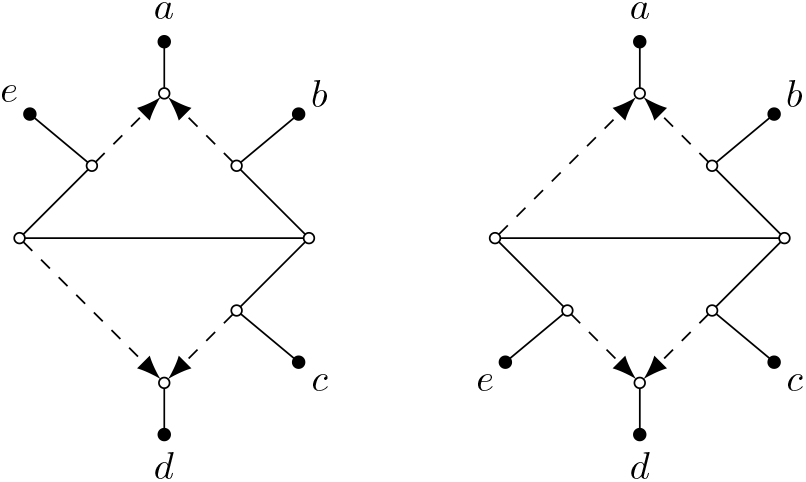
Two distinct simple, level-2 quinnets on leaf set *S* = {*a, b, c, d*, e} that are distinguishable by Lemma 2.14d.

### 2.4. Distinguishing Three-Leaf Trees from Trinets

In this section, we describe an inequality for the JC model that allows us to distinguish a three-leaf tree from a strict level-1 or level-2 trinet (as elsewhere in the paper, all parameters are assumed to belong to (0, 1)). This inequality helps us prove distinguishability results for many pairs of level-2 networks, since many level-2 networks have induced subnetworks that are trees.

Recall that under the JC model, for every edge *e* we have 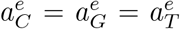, and 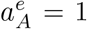. It follows that many Fourier coordinates coincide. For the Fourier coordinates *q* under the JC model for a trinet, we therefore write *q*_110_ as shorthand for any of the values *q*_*CCA*_ = *q*_*GGA*_ = *q*_*TTA*_, since all of these values are the same under the JC model. We define *q*_101_ and *q*_011_ similarly. We write *q*_111_ to denote any of *q*_*CGT*_ = *q*_*CTG*_ = *q*_*GCT*_ = *q*_*GTC*_ = *q*_*TGC*_ = *q*_*TCG*_, as again all of these values are the same under the JC model.

#### Proposition 2.15

*Consider the Fourier coordinates q associated to a binary, semi-directed level-2 trinet N under the JC model. Then the invariant*

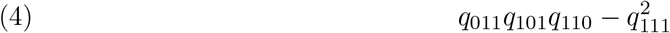

*evaluates to zero if N is a three-leaf tree and is strictly positive if N is a strict level-1 or strict level-2 trinet*.

*Proof*. Throughout the proof, we simplify the notation for the Fourier parameters as follows. In the description in Section 2.1.2, the parameters were notated as 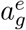, where *e* is an edge and *g* is a group element. However, in the JC model, we have that 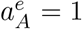, and 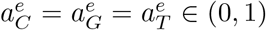.

Hence, we use a single letter to denote the parameter value 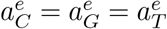, and we use a different letter *a, b, c*, … for each edge in the network.

First consider the case where *N* is a 3-leaf tree. Then using the JC Fourier parameters shown in Figure 7, *q*_111_ = *abc, q*_110_ = *ab, q*_101_ = *ac*, and *q*_011_ = *bc*, so the expression of (4) evaluates to 0.

**Figure 7.**
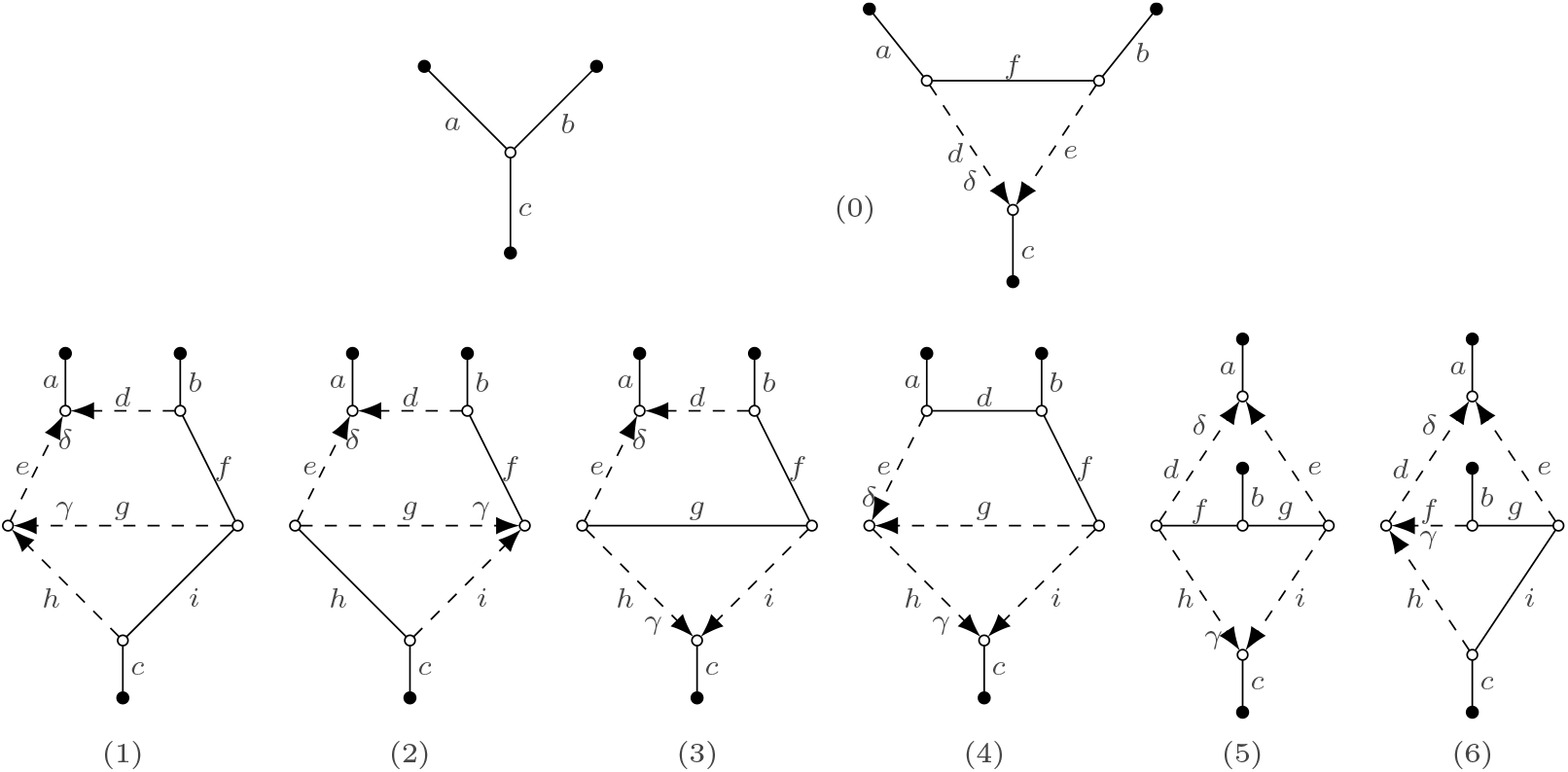
All eight simple level-2 trinets up to labeling the leaves, with *a, b*, …, *i* denoting edge labels and δ and *γ* denoting inheritance probabilities. The edge and trinet labels are used in the proof of Proposition 2.15.

Note that if *N* is not a 3-leaf tree, it can either be the strict level-1 trinet, or one of the six strict level-2 trinets (see Figure 7 for their numbering). We now show that the invariant is strictly positive if *N* is the trinet of type 0 and defer the very similar calculations for the trinets of types 1-6 to Appendix A.

#### Trinet 0

Let *a*-*f* be the Fourier parameters associated to the edges of *N* (see also Figure 7). Furthermore, let δ ∈ (0, 1) be the inheritance probability. Then, *N* has the following Fourier parameterizations under the JC model:

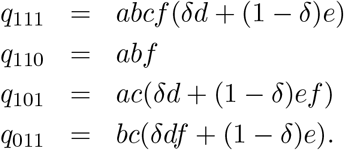

Thus,

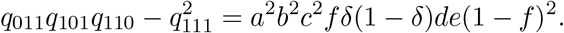

Since all parameters are in (0, 1), 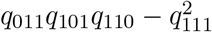 is strictly positive.

While we have only been able to find a proof of Proposition 2.15 for level-2 networks, and only by analyzing each network on a case by case basis, we conjecture that the analog of this result should hold for trinets of arbitrary level (as always assuming that there are no 2-blobs).

#### Conjecture 2.16

*Let N be a strict level-k, semi-directed trinet with k >* 0. *Then, under the JC model, the invariant* (4) *is strictly positive*.

We call a leaf *x* of a simple quarnet *N* on *S* a *prune leaf*, if *N* | _*S \* {*x*}_ is a three-leaf tree. As an example, the leaves in the top left of the planar representations in Figure 5 of the quarnets of type 0, 1c, 1d, 1e, 2d, and 3b are prune leaves. As a direct application of Proposition 2.15 and the fact that the quarnets of type 0, 1c, 1d, 1e, 1f, 2d, 3b and 3e have exactly one prune leaf, and the other quarnets have no prune leaves, we obtain the following useful result, which will be used to prove our main theorem.

#### Corollary 2.17

*Let N*_1_ *and N*_2_ *be two binary level-2 semi-directed phylogenetic networks on the same leaf set X. Let S be a 4 leaf subset of X. Then, under the JC model:*

a. *if N*_1_ | _*S*_ *is a type 0, 1c, 1d, 1e, 1f, 2d, 3b or 3e quarnet and N*_2 *S*_ *is a type 1a, 1b, 2a, 2b, 2c, 3a, 3c or 3d quarnet, then N*_1_ *and N*_2_ *are distinguishable;*
b. *if N*_1_ | _*S*_ *and N*_2_ | _*S*_ *are distinct quarnets of type 0, 1c, 1d, 1e, 1f, 2d, 3b or 3e with different prune leaves, then N*_1_ *and N*_2_ *are distinguishable*.

## 3. RESULTS

### 3.1. Identifiability of Level-1 Networks

Building on the results of the previous subsections we are now ready to prove the main identifiability result for level-2 networks. Before we do this, we first strengthen the best existing identifiability results for semi-directed level-1 networks under a network-based Markov model from Gross et al. (2021). They proved that the network parameter of a JC, K2P or K3P network model is generically identifiable with respect to the class of *n*-leaf triangle-free, level-1 semi-directed networks with a fixed number of *r* ≥ 0 reticulation nodes. Using our previous results, we are able to considerably strengthen these results in the case of JC and K2P constraints. In particular, the following theorem shows that not only can we identify such level-1 networks if they have the same number of reticulation nodes, but also in the case where the number of reticulation nodes differs. Furthermore, for the JC constraints we can relax the restriction that the networks need to be triangle-free.

**Theorem 3.1**

a. *The network parameter of a JC network model is generically identifiable up to the choice of reticulation nodes in triangles, with respect to the class of models where the network parameter is an n-leaf binary level-1 semi-directed phylogenetic network*.
b. *The network parameter of a K2P network model is generically identifiable with respect to the class of models where the network parameter is an n-leaf binary, triangle-free, level-1 semi-directed phylogenetic network*.

*Proof*. The tree-of-blobs of a network is generically identifiable under both the JC and K2P constraints by Corollary 2.12. By Proposition 2.15, triangles can also be distinguished from trivial blobs under the JC constraints. The results then follow from the fact that simple level-1 networks (or, *sunlets*) with at least four leaves are identifiable under the JC (Gross and Long, 2018) and K2P (Hollering and Sullivant, 2021) constraints.

### 3.2. Identifiability of Level-2 Networks

Piecing together distinguishability results from previous sections, Lemmas A.2 and A.3 in Appendix A.4 show distinguishability under the JC constraints of classes of outer-labeled planar and not outer-labeled planar simple level-2 networks, respectively. These lemmas now allow us to prove our main result in the next theorem. Note that Theorem 3.1a is not a special case of this theorem, since here we also assume that the network is triangle-free.

#### Theorem 3.2

*The network parameter of a JC network model is generically identifiable with respect to the class of models where the network parameter is an n-leaf binary, triangle-free, strongly tree-child, level-2 semi-directed phylogenetic network*.

*Proof*. Let *N*_1_ and *N*_2_ be two distinct *n*-leaf binary, triangle-free, strongly tree-child, level-2 semi-directed phylogenetic networks on the same leaf set *X*. If *N*_1_ and *N*_2_ have non-isomorphic trees-of-blobs, they are distinguishable by Corollary 2.12. Since a network is strongly tree-child if and only if the simple networks induced by each blob are strongly tree-child — a fact that follows from Theorem 5 in Holtgrefe et al. (2026), see also Definition 2.3 — we may consider each blob separately and it will be enough to assume that *N*_1_ and *N*_2_ are both simple networks. Note that *N*_1_ and *N*_2_ cannot contain nontrivial 3-blobs since those are not triangle-free or not strongly tree-child (see Figure 7), so we may also assume *N*_1_ and *N*_2_ have at least 4 leaves.

If *N*_1_ and *N*_2_ are both not outer-labeled planar, then the strongly tree-child assumption forces both networks to be stack-free. Then, they are distinguishable by Lemma A.2. If *N*_1_ and *N*_2_ are both outer-labeled planar, they are distinguishable by Lemma A.3. Lastly, without loss of generality, assume that *N*_1_ is outer-labeled planar and *N*_2_ is not. Then, *N*_2_ is strict level-2 and there is some *S* ⊆ *X* with |*S*| = 4 such that the quarnet *N*_2_ | _*S*_ is stack-free and not outer-labeled planar and thus is of type 3a, 3b, 3c or 3d. On the other hand, *N*_1 *S*_ must be outer-labeled planar and can thus not be of type 3a, 3b, 3c or 3d. But the type 3a, 3b, 3c or 3d quarnets are the only level-2 quarnets that display exactly three quartet trees. Therefore, *N*_1_ and *N*_2_ are distinguishable by Theorem 2.11, which lets us distinguish quarnets on the basis of their displayed trees.

### 3.3. Identifiability Beyond Level-2

In Section 2.2 we proved that semi-directed networks with different displayed quartets are identifiable (Theorem 2.11). Consequently, we also obtained two interesting identifiability results regarding the tree-of-blobs (Corollary 2.12) and the congruent circular orderings of networks (Corollary 2.13), which we later used to obtain our proof of identifiability for level-2 networks. We summarize these three results here because they are also of independent interest (see e.g. Allman et al. (2023); Rhodes et al. (2025) for similar results under coalescent based models), especially because the results pertain to networks of any level. Note that these results hold under both the JC and K2P models and under the assumption that inheritance probabilities and branch lengths are finite and nonzero.

#### Theorem 3.3

*Let N*_1_ *and N*_2_ *be two binary semi-directed phylogenetic networks on the same leaf set X. Let* 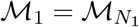 *and* 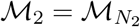 *be the associated JC or K2P network models*.

a. *If there is a* 4 *element subset S* ⊆ *X such that the quarnets N*_1_ | _*S*_ *and N*_2_ | _*S*_ *have different sets of displayed quartets, then M*_1_ ∩ *M*_2_ = ∅.
b. *If the trees-of-blobs of N*_1_ *and N*_2_ *are not isomorphic, then M*_1_ ∩ *M*_2_ = ∅.
c. *If N*_1_ *and N*_2_ *are both outer-labeled planar and they have different sets of congruent circular orders, then M*_1_ ∩ *M*_2_ = ∅.

## 4. Discussion

We have shown that the class of strongly tree-child, triangle-free, level-2, binary semi-directed networks is identifiable under the JC model of evolution. Our results indicate that, under this model, nucleotide sequence data provides enough information to infer semi-directed networks from this class, which can be used as a realistic estimate of a phylogeny from DNA sequence data. This is the first identifiability result of a large class of networks beyond the level-1 framework. As detailed by Kong et al. (2022), a tree-child scenario is more likely to appear in nature than a non-tree-child scenario, since hybridization is considered to be an uncommon evolutionary event, further supporting the biological relevance of our identified class.

The results in this paper are not limited to level-2 networks. Specifically, we have also shown that the tree-of-blobs of any binary semi-directed network and the circular ordering of the subnetwork around each outer-labeled planar blob are identifiable under both the JC and K2P models, thereby extending existing results in Allman et al. (2023); Rhodes et al. (2025) to other classes of models. Identifiability of the tree-of-blobs also allowed us to considerably strengthen previous identifiability results for level-1 networks under Markov models (Gross et al., 2021).

It remains an open problem to try to prove identifiability of level-2 (and higher level) networks for broader classes of networks than we have shown in this paper. However, there are already challenges such that for the Markov models it is not possible to identify a network structure in full generality. For example, stacked reticulation nodes always lead to non-identifiability issues, as shown in Sullivant (2025). Even for level-1, it remains open to prove identifiability results for nonbinary networks.

The combinatorial component of our proof, which relies on algebraic distinguishability proofs for small networks, may provide a blueprint to prove similar level-2 identifiability results under other models of evolution. For instance, to lift our level-2 identifiability result to the K2P model, it would suffice to generalize Lemma 2.14 and Proposition 2.15, which establish distinguishability for specific combinations of small networks. Similarly, the combinatorial part of our proofs could be useful when attempting to generalize our results to settings with incomplete lineage sorting. Going beyond the level-2 assumption will likely require further improvements to the existing algebraic tools. Notably, it is already known that semi-directed level-3 networks are not encoded by their quarnets (Huber et al., 2025). As a result, achieving identifiability of (a subclass of) level3 networks may necessitate analyzing numerous combinations of induced level-3 quinnets (5-leaf networks), leading to a combinatorial explosion of combinations compared to level-2 quarnets.

On the algebraic side, there are many open questions related to level-2 networks, and even quarnets. For example, while we computed some phylogenetic invariants for these networks in some cases, in most cases it remains open to determine the vanishing ideal of the model, or other basic algebraic properties of the models. We found instances of pairs of networks that have the same underlying algebraic matroid but different vanishing ideals. There were examples of networks that had a lower dimension than would be expected based on a simple parameter count. Developing tools to explain these phenomena leads to interesting algebraic challenges, and may provide new tools for proving deeper identifiability results.

Finally, on the algorithmic side, our findings open several promising research directions. Previous identifiability proofs for level-1 networks under the models considered here (Gross et al., 2021) have resulted in theoretical algorithms that puzzle together quarnets to construct a complete level-1 network (Huebler et al., 2025; Frohn et al., 2025), ultimately contributing to a quarnet-based practical tool for inference of level-1 networks (Holtgrefe et al., 2025b). Building on our results, similar theoretical algorithms could serve as a foundation for consistent level-2 network inference methods. However, such algorithms, which rely on assembling multiple quarnets, also require consistent methods for inferring these quarnets from biological data. Two existing approaches, both based on invariants, have been developed for level-1 quarnets (Barton et al., 2026; Martin et al., 2025). To establish a complete pipeline from sequence data to level-2 network reconstruction, analogous techniques may be necessary for level-2 quarnets.

## Funding Acknowledgment

This material is based upon work supported by the National Science Foundation under Grant No. DMS-1929284 while the authors were in residence at the Institute for Computational and Experimental Research in Mathematics in Providence, RI, during the semester program on “Theory, Methods, and Applications of Quantitative Phylogenomics”. AE was supported by the Wisconsin Alumni Research Foundation, the Georgia Benkart Legacy Fund and NSF Award DMS-2023239. EG was supported by the National Science Foundation under Grant No. DMS-1945584. MF, NH, LvI & MJ were supported by the Dutch Research Council (NWO) under Grant No. OCENW.M.21.306, and LvI & MJ also under Grant No. OCENW.KLEIN.125.

## Acknowledgment

We thank David Morrison for helpful comments and suggestions to improve the readability of the paper for a biological audience.

## Data Availability Statement

The `Macaulay2` scripts necessary to reproduce the proof of Lemma 2.14 and the `Maple` scripts used to verify the proof of Proposition 2.15 can be accessed at: https://doi.org/10.5281/zenodo.20193239.

## Appendix Omitted proofs

Our main result relies on a mixture of algebraic, computational and combinatorial results, with several of the longer proofs presented in this appendix.

### A.1. Distinguishing Networks with Different Displayed Quartets

#### Proposition 2.10

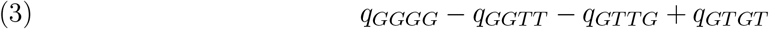

*Proof*. Under the tree 12|34 we have

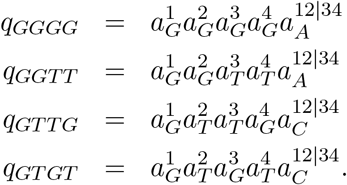

Since 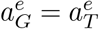 for all edges *e*, we see that *q*_*GGGG*_ = *q*_*GGTT*_ and *q*_*GTTG*_ = *q*_*GTGT*_, so expression 3 evaluates to 0.

Similarly, under the tree 14|23 we have

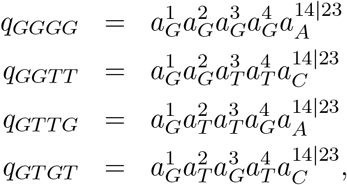

so *q*_*GGGG*_ = *q*_*GTTG*_ and *q*_*GGTT*_ = *q*_*GTGT*_. Thus, expression 3 again evaluates to 0.

On the other hand, under the tree 13|24 we have that

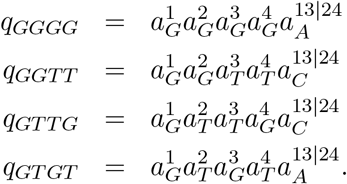

So, expression 3 becomes 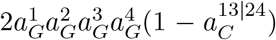 since 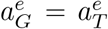 and 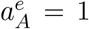 for all edges *e*.

Because 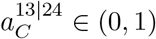, this shows that expression 3 is positive.

The following result is presented in a slightly different form in Rhodes et al. (2025). Since the original source presents only an implicit proof in a different context (Allman et al., 2023), we prove part (a) here as well.

#### Proposition A.1

*Let N be a binary semi-directed phylogenetic network on 4 leaves {a, b, c, d}. Then*,

a. *N induces the nontrivial split ab* | *cd if and only if N displays solely the quartet with nontrivial split ab cd; if N is simple and outer-labeled planar (congruent with circular order* (*a, b, c, d*)*), then N*
b. *displays exactly the two quartets with nontrivial splits ab*|*cd and ad*|*bc*.

*Proof of part (a)*. The only-if direction of part (a) is trivial since every quartet displayed by *N* has the cut-edge induced by the split *ab* | *cd* in its embedding. Thus, all the quartets displayed by *N* have the split *ab* | *cd*.

For the if-direction of part (a), we use induction on the level *k* of the network. Assume that *N* does not induce any nontrivial split. For *k* = 0 this is not possible and for *k* = 1, *N* will contain a 4-cycle, causing it to display two quartets with different splits, so we are done. For *k* ≥ 2, pick a reticulation *r* with a leaf *x* below it. Create two level-(*k* − 1) networks *N*_1_, *N*_2_ by deleting one of the two reticulation edges entering *r*, undirecting the other reticulation edge and exhaustively deleting reticulation nodes with in-degree 2 and out-degree 0, suppressing non-reticulate degree-2 nodes and suppressing parallel edges. If at least one of *N*_1_, *N*_2_ does not induce a nontrivial split, then it displays at least two quartets by induction. Hence, *N* also displays at least two quartets. Now consider the remaining case, that both *N*_1_ and *N*_2_ induce a nontrivial split. Then it is sufficient to show that *N*_1_ and *N*_2_ display different nontrivial splits.

Suppose that *N*_1_ and *N*_2_ display the same nontrivial split, say *xy wz*. Let *N*′ be the mixed graph obtained from *N* by deleting *x* and exhaustively deleting resulting reticulation nodes with in-degree 2 and out-degree 0. Note that *N*′ is connected since otherwise *N* could not be oriented as a directed phylogenetic network. Let *e* be the cut-edge of *N*′ inducing the split *xy* | *wz* with maximum distance from *y*. Let *N*^*′′*^ be the connected component of the mixed graph obtained from *N*′ by deleting *e*, such that *N*^*′′*^ contains *w* and *z*. Since *N* does not have a cut-edge inducing the split *xy* | *wz*, it contains a directed path from *N*^*′′*^ to the neighbour of *x*, which does not use *e*. Hence, *N*_*i*_ contains a directed path from *N*^*′′*^ to the neighbour of *x* for some *i* ∈ {1, 2} that does not use *e*. However, this implies that *N*_*i*_ does not have a cut-edge inducing the split *xy* | *wz*, a contradiction.

### A.2. Distinguishing Small Networks using Invariants and Matroids

#### Lemma 2.14

*Remainder of the proof*

a. This computation is verified in the file `Lemma214a.m2`. First of all, we only need to look at the different labelings of the leaves of quarnet 1b that have the same displayed trees. With the leaves labeled clockwise starting in the upper left, this amounts to looking at the eight circular orders

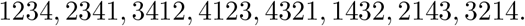

Here we want to show that each of these circular orders is distinguished from the others except in the case that the top middle leaf gets the same label, e.g. we do not need to distinguish the pair 1234, 3214. In fact, separate computations show that those pairs have the same ideal. As explained below Proposition 2.6, this does not prove that the corresponding models are indistinguishable, although it does provide evidence in favor of this conclusion. For each of the pairs we want to distinguish, there is a subset of the variables that has the property that the corresponding columns of the Jacobian matrices under the networks have different ranks. These subsets are listed in “successList” in the file. For example, for the first pair (corresponding to the two orders 1234, 2341, the set of columns of the Jacobian matrix

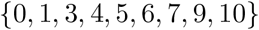

yield different ranks in the two networks. These correspond to the indeterminates

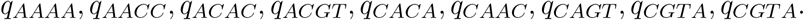

Running the code and producing the term “listofrankpairs” shows the ranks of the corresponding submatrices of the Jacobians for these two networks. It has rank 8 for ordering 1234 and rank 9 for ordering 2341.
b. The proof of (b) follows a similar strategy as (a), and the corresponding computation is in the file `Lemma214b.m2`.
c. The proof of (c) is established by finding phylogenetic invariants that distinguish each pair of networks. In the file `InvariantForThreeA.m2`, we give a degree 6 invariant with 53 terms that distinguishes quarnet 3a with the labeling 1234 from all other labeled 3a, 3b, and 3d quarnets. In the file `InvariantForThreeC.m2`, we give a degree 5 invariant with 34 terms that distinguishes the 3c quarnet with the labeling 1234 from all other labeled 3a, 3b, and 3d quarnets, and in the file `InvariantForThreeD.m2`, we give a degree 5 invariant with 50 terms that distinguishes the 3d quarnet with labeling 1234 from all other labeled 3a, 3b, and 3d quarnets. The computation verifying the statements is in the file `Lemma214c.m2`.
d. The proof of (d) follows a similar strategy as (a), and the corresponding computation is in the file `Lemma214d.m2`. In this case there is just one pair of networks to compare. Considering the set of indeterminates

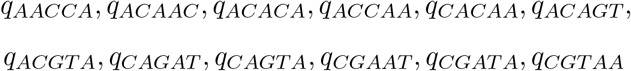

the code shows that for the left network, the corresponding rank of the submatrix of the Jacobian is 12, and for the right network it is 11.

### A.3. Distinguishing Three-Leaf Trees from Trinets

#### Proposition 2.15

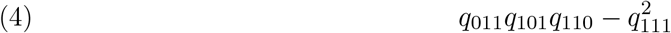

*Remainder of the proof*. In the main text we showed that the invariant (4) evaluates to zero for the three-leaf tree, and that it is positive on the trinet of type 0 (see Figure 7 for the trinet numbering), assuming the parameter values are in (0, 1). Here, we complete the proof by providing the Fourier parameterizations of the trinets 1-6 under the JC model (using the parameters as given in Figure 7) and proving positivity of expression (4) by providing a suitable decomposition. Correctness of the decompositions can be verified using the `Maple` files `trinet1.mw` – `trinet6.mw`.

#### Trinet 1

As depicted in Figure 7, let *a*-*i* be the nontrivial Fourier parameters associated to the edges of *N*. Furthermore, let δ and *γ* be the two inheritance probabilities. Under the JC model, *N* has the following Fourier parameterizations:

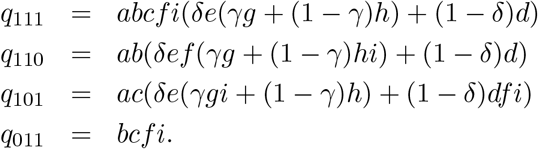

We now compute *q*_110_*q*_101_*q*_011_ − *q*^2^ and look at the coefficients for δ^2^, (1 − δ)^2^ and δ(1 − δ).

The coefficient for δ^2^ is

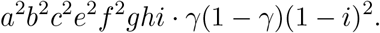

The coefficient for (1 − δ)^2^ is as follows:

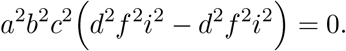

Lastly, the coefficient for (1 − δ)δ becomes

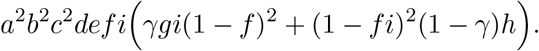

Since the parameter values are in (0, 1), the coefficients for δ^2^ and (1 − δ)δ are positive, while the coefficient for (1 − δ)^2^ was equal to zero. All in all, this shows that the invariant 4 is positive.

#### Trinet 2

The Fourier parameterizations of Trinet 2 are as follows:

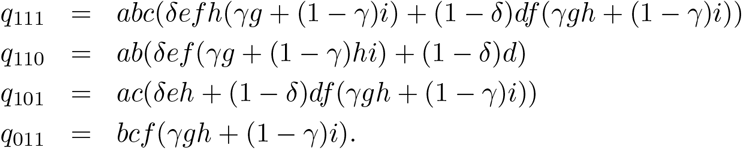

For the invariant we get that

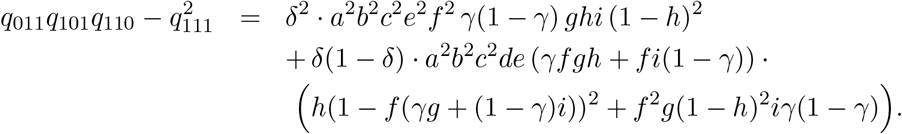

which is positive since all parameter values are in (0, 1).

#### Trinet 3

The Fourier parameterizations for Trinet 3 are

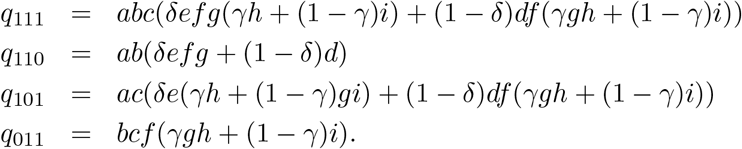

Then, one obtains that

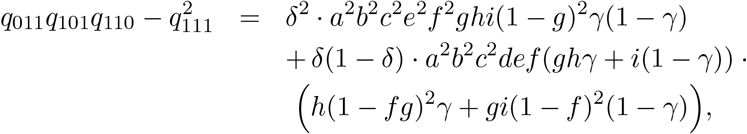

which is again positive since all parameter values are in (0, 1).

#### Trinet 4

The Fourier coordinates of Trinet 4 are:

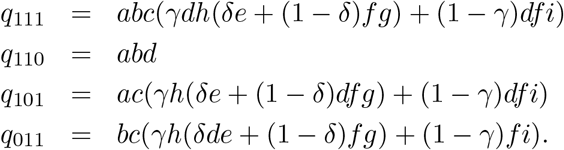

The invariant for Trinet 4 is

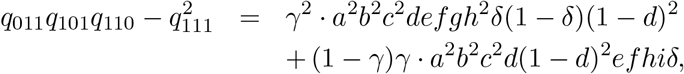

which is positive under our parameter constraints.

#### Trinet 5

The Fourier coordinates are now as follows:

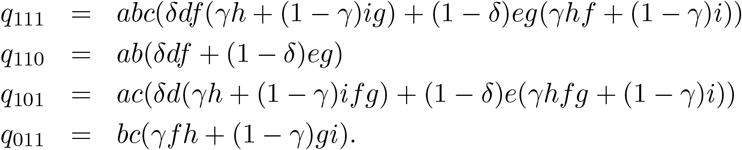

The invariant then becomes

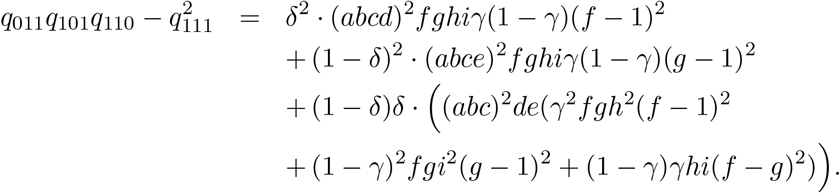

This can again be seen to be positive since all parameter values are in (0, 1).

#### Trinet 6

The Fourier parameterizations of Trinet 6 are

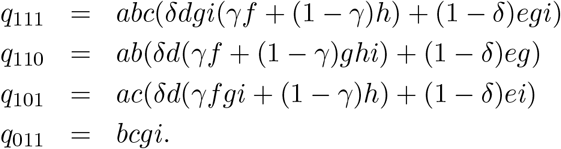

After simplification, the invariant becomes

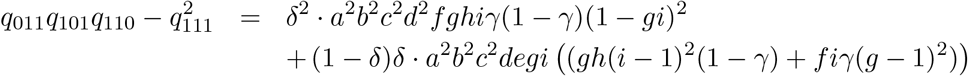

This is positive provided all parameters are positive and smaller than 1.

### A.4. Distinguishing Large Level-2 Networks

This section focuses on distinguishing certain pairs of *n*-leaf, simple, semi-directed, level-2 networks under the JC model. We state our results in the form of two key lemmas that rely on our previous results to argue that any pair of considered networks has a subnetwork that distinguishes them. We implicitly use Proposition 2.8 to then extend the distinguishability to the full network.

#### Lemma A.2

*Let N*_1_ *and N*_2_ *be distinct binary, stack-free, simple, semi-directed level-2 phylogenetic networks on the same leaf set. If neither N*_1_ *nor N*_2_ *is outer-labeled planar, then N*_1_ *and N*_2_ *are distinguishable under the JC constraints*.

*Proof*. Let *X* be the leaf set of the two networks. First note that both *N*_1_ and *N*_2_ are strict level-2, otherwise they would be outer-labeled planar. The only strict level-2 quarnets that are stack-free and not outer-labeled planar are of type 3a, 3c or 3d (see Figure 5 for the quarnet numbering). Hence, if | *X* |= 4, *N*_1_ and *N*_2_ are distinguishable by Lemma 2.14c and we may thus assume that | *X* |≥ 5.

Let *G* be the *generator* of *N*_1_, i.e., the mixed multi-graph obtained from *N*_1_ by deleting all leaves and suppressing degree-2 nodes. Similarly, we let *H* be the generator of *N*_2_. By Lemma 4.2 in Huber et al. (2025), there are exactly two possibilities for *G* and *H*, depicted in Figure 8: the generators Γ_1_ and Γ_2_. The edges and reticulation nodes with in-degree 2 and out-degree 0 of Γ_1_ and Γ_2_ are called *sides*. For a side *S* of Γ_1_ or Γ_2_, we call a leaf *s on side S* if leaf *s* is attached to side *S* in the original network. We now consider three cases. We will implicitly use that for every *S* ⊆ *X* with | *S* | = 4 the two quarnets *N*_1 *S*_ and *N*_2 *S*_ either both have the same nontrivial split or neither has a nontrivial split, otherwise the two quarnets have a distinct tree-of-blobs and the networks are already distinguishable by Corollary 2.12.

**Figure 8.**
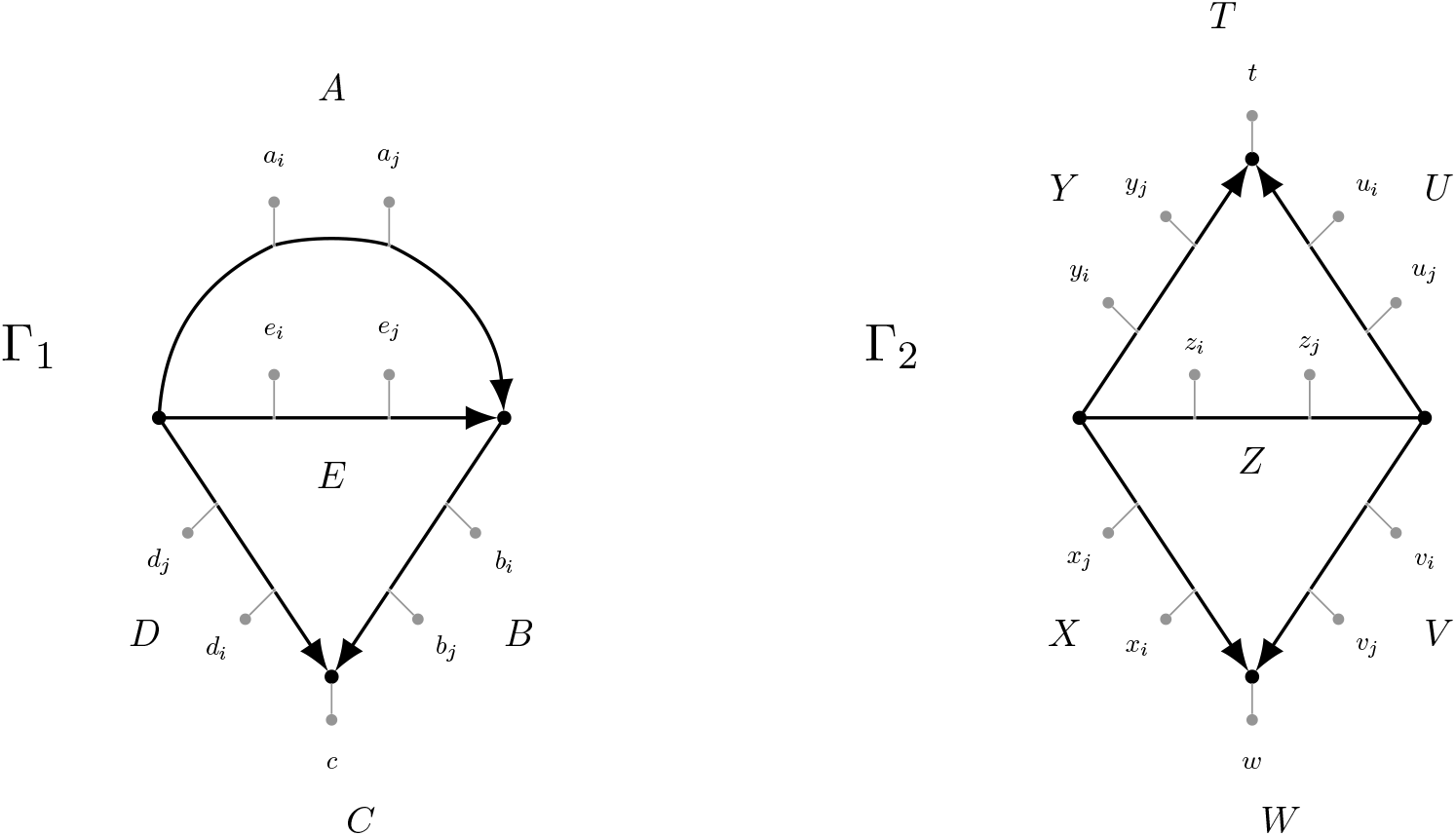
The two generators Γ_1_ and Γ_2_ (in thick black) of a simple, strict level-2, semi-directed network (Huber et al., 2025, Lem. 4.2). The sides {*A, B, C, D, E*} and {*T, U, V, W, X, Y, Z*} of the two generators are the places where the leaves (in gray) might be attached in the corresponding semi-directed network.

#### Case 1

*G* ≇ *H*. Without loss of generality we assume that *G* ≅ Γ_1_ and *H* ≅ Γ_2_. In *N*_1_, let *a* be any leaf on side *A, b* any leaf on side *B, c* the leaf on side *C*, and *e* any leaf on side *E*. Note that *a* and *e* exist since *N*_1_ is not outer-labeled planar, and that *b* exists since *N*_1_ is stack-free. Then, the quarnet *N*_1_ | _*S*_ with *S* = *a, b, c, e* is of type 3a. Now observe that quarnets of type 3a, 3b and 3e each contain one reticulation node whose child is not a leaf. Since only leaves can be added to sides *T* and *W* in Γ_2_, this means *N*_2 *S*_ cannot be of type 3a, 3b or 3e. Thus, *N*_2_ | _*S*_ is distinct from *N*_1 *S*_. Specifically, it is of type 0, 1, 2, 3c or 3d. Then, if *N*_2 *S*_ is of type 0, 1 or 2, *N*_1_ and *N*_2_ are distinguishable by Theorem 2.11, since *N*_2_ | _*S*_ then displays at most two quartet trees, whereas *N*_1_ | _*S*_ displays three quartet trees. Otherwise, if *N*_2_ | _*S*_ is a quarnet of type 3c or 3d, *N*_1_ and *N*_2_ are distinguishable by Lemma 2.14c.

#### Case 2

*G* ≅ *H* ≅ Γ_1_. Let *S* = {*a, b, c, e*} be an arbitrary set of leaves as in Case 1, such that *N*_1_ | _*S*_ is a quarnet of type 3a. Similar reasoning as in Case 1 then shows that *N*_1_ and *N*_2_ are distinguishable if for some choice of *S* the quarnet *N*_1_ | _*S*_ is distinct from *N*_2_ | _*S*_. Hence, we may assume that *N*_1_ | _*S*_ and *N*_2_ | _*S*_ are isomorphic for any valid choice of *S*. It follows that *N*_1_ and *N*_2_ share the same leaves on the sides *A, B, C* and *E* of their generator (modulo swapping sides *A* and *E*), and therefore also on side *D*. Then, since by assumption *N*_1_ and *N*_2_ are distinct and |*X* | ≥ 5, we have that for some side Σ ∈ *{A, B, D, E}* (with |Σ| ≥ 2) two leaves *σ*_*i*_ ∈ Σ and *σ*_*j*_ ∈ Σ are ordered differently in the two networks. Now, let *c* be the leaf on side *C* of *N*_1_ and *N*_2_. First suppose that Σ ∈ *{A, D, E}* and let *b* be a leaf on side *B* of *N*_1_ and *N*_2_ (which exists since the networks are stack-free). Then, the quarnets *N*_1_ | _*S*_ and *N*_2_ | _*S*_ with *S* = {*σ*_*i*_, *σ*_*j*_, *b, c*} are both simple and outer-labeled planar, i.e. both are congruent with a single circular ordering of *S*. But since *σ*_*i*_ and *σ*_*j*_ are ordered differently on side Σ in the two networks, these circular orders cannot be the same. Hence, *N*_1_ and *N*_2_ are distinguishable by Corollary 2.13, which states that two simple and outer-labeled planar quarnets are distinguishable if they are not congruent with the same circular ordering. The case where Σ = *B* follows a similar argument, instead considering the subset of leaves *S* = {*σ*_*i*_, *σ*_*j*_, *a, c*}, where *a* is a leaf from side *A* of both networks (which exists since the networks are not outer-labeled planar).

#### Case 3

*G* ≅ *H* ≅ Γ_2_. In *N*_1_, let *t* be the leaf on side *T, w* the leaf on side *W, z* any leaf on side *Z*, and *σ* any other leaf. Note that *t, w* and *z* exist since *N*_1_ is not outer-labeled planar. Then, the quarnet *N*_1 *S*_ with *S* = {*t, w, z, σ*} is of type 3c or 3d. If for some valid choice of *S* the quarnet *N*_2_ | _*S*_ is of type 0, 1 or 2, we get that *N*_1_ and *N*_2_ are distinguishable by Theorem 2.11, since *N*_2_ | _*S*_ then displays at most two quartet trees, whereas *N*_1 *S*_ displays three quartet trees. Now note that if *N*_2 *S*_ is not of type 0, 1 or 2, it can only be of type 3c or 3d. By Lemma 2.14c, two distinct quarnets of type 3c or 3d are distinguishable, so we may now assume that *N*_1_ | _*S*_ and *N*_2_ | _*S*_ are isomorphic for all valid choices of *S*. It follows that *N*_1_ and *N*_2_ share the same leaves on the sides *T, Y ∪ U, Z, X ∪ V* and *W* of their generators (modulo swapping *Y, T, U* with *X, W, V*).

If | *Y* ∪ *U* ∪ *X* ∪ *V* | ≤ 1 for *N*_1_ and *N*_2_, then *N*_1_ and *N*_2_ share the same leaves on every side (modulo swapping *Y* with *U* and swapping *X* with *V*). If | *Y* ∪ *U* ∪ *X* ∪ *V* | ≥ 2 for *N*_1_ and *N*_2_, let *σ* and *π* be two arbitrary leaves in this set. Furthermore, let *t* be the leaf on side *T* of *N*_1_ and *N*_2_ and *w* the leaf on side *W* of *N*_1_ and *N*_2_. Then, *N*_1 *S*_ and *N*_2 *S*_ with *S* = {*σ, π, t, w*} are both simple and outer-labeled planar quarnets, i.e. both congruent with a single circular order of *S*. If for some valid choice of *S* the induced orderings are not the same, then *N*_1_ and *N*_2_ are distinguishable by Corollary 2.13. Thus, we may assume that these circular orders are the same for any such set *S*, again implying that *N*_1_ and *N*_2_ share the same leaves on every side (modulo swapping *Y* with *U* and swapping *X* with *V*). Furthermore, the leaves on these four sides must be ordered the same.

The only remaining way for *N*_1_ and *N*_2_ to be distinct is then that there are two leaves *z*_*i*_ and *z*_*j*_ on side *Z* of *N*_1_ and *N*_2_ that are not ordered the same in the two networks. Let *σ* be a leaf unequal to *z*_*i*_ or *z*_*j*_ from some side Σ ∈ {*U, V, X, Y, Z}* (which exists since |*X* | ≥ 5). If Σ ∈ {*U, Y, Z}* then we consider the set *S* = {*σ, t, z*_*i*_, *z*_*j}*_ with *t* the leaf on side *T* of both networks. Otherwise, if Σ *V, X*, we instead pick the other leaf below the reticulation *w* (on side *W* of both networks) and consider the set *S* = *σ, w, z*_*i*_, *z*_*j*_. In both cases, the quarnets *N*_1 *S*_ and *N*_2 *S*_ are of type 0 (i.e. they are level-1 quarnets with a 4-cycle) and they induce a unique circular ordering of *S*. But since *z*_*i*_ and *z*_*j*_ are ordered differently on side *Z* in the two networks, these induced orders cannot be the same. Hence, *N*_1_ and *N*_2_ are distinguishable by Corollary 2.13.

#### Lemma A.3

*Let N*_1_ *and N*_2_ *be distinct binary, triangle-free, strongly tree-child, simple, semi-directed level-2 phylogenetic networks on the same leaf set. If N*_1_ *and N*_2_ *are both outer-labeled planar, then N*_1_ *and N*_2_ *are distinguishable under the JC constraints*.

*Proof*. Let be the leaf set of the two networks. If *N*_1_ and *N*_2_ are both level-1 networks, they are distinguishable by Gross and Long (2018). If exactly one of them (say *N*_1_) is a strict level-2 network, then there will be some *S* ⊆ *X* with | *S* | = 4 such that *N*_1 *S*_ is a type 1b or 2a quarnet, since *N*_1_ is triangle-free, strongly tree-child and outer-labeled planar (see Figure 5 for the quarnet numbering). Because *N*_2_ is level-1, *N*_2_|_*S*_ is either a type 0 quarnet or a quarnet inducing a nontrivial split. In the first case, *N*_1_|_*S*_ and *N*_2_|_*S*_ are distinguishable by Corollary 2.17a, whereas in the second case the two quarnets have a distinct tree-of-blobs and are distinguishable by Corollary 2.12. Thus, we may assume that *N*_1_ and *N*_2_ are both strict level-2. Now note that the only outer-labeled planar strict level-2 quarnets that are strongly tree-child and triangle-free are of type 2a. Hence, if |*X* | = 4, the two networks are distinguishable by Lemma 2.14b. Thus, we will also assume that |*X* | ≥ 5.

As in the proof of Lemma A.2, we let *G* be the generator of *N*_1_ and *H* the generator of *N*_2_. See again Figure 8 for the two possible generators Γ_1_ and Γ_2_. Since *N*_1_ and *N*_2_ are outer-labeled planar, we may assume in any case that the no leaves are attached to side *E* of Γ_1_ and to side *Z* of Γ_2_. We again consider three cases. Note that no quarnets of *N*_1_ or *N*_2_ can be of type 3, since both networks are outer-labeled planar. As in the proof of the previous lemma, we will implicitly use that for every *S*⊆ *X* the two quarnets *N*_1_ | _*S*_ and *N*_2_ | _*S*_ with |*S*| = 4, either both have the same nontrivial split or neither has a nontrivial split, otherwise the two quarnets have a distinct tree-of-blobs and the networks are already distinguishable by Corollary 2.12. Furthermore, by Corollary 2.13, we can assume that *N*_1_ and *N*_2_ share the same circular ordering of their leaf set *X*, otherwise the two networks are again distinguishable.

**Case 1:** *G* ≇ *H*. Without loss of generality we assume that *G* ≅ Γ_1_ and *H* ≅ Γ_2_. In *N*_1_, let {*a*_*i*_, *a*_*j*_} be any pair of leaves on side *A, c* the leaf on side *C* and *d* any leaf on side *D*. Note that *a*_*i*_ and *a*_*j*_ exist since *N*_1_ is triangle-free and that *d* exists since *N*_1_ is strongly tree-child. Then, the quarnet *N*_1 *S*_ with *S* = {*a*_*i*_, *a*_*j*_, *c, d}* is of type 2d with *c* being its prune leaf (see the definition before Corollary 2.17). If *N*_2_ | _*S*_ is of type 1a, 1b, 2a, 2b or 2c, the two quarnets are distinguishable by Corollary 2.17a, whereas the two quarnets are distinguishable by Corollary 2.17b if *N*_2_ | _*S*_ is of type 0, 1c, 1d, 1e, 1f or 2d with a prune leaf *σ* ≠ *c*. It follows that we may assume that *N*_2_ | _*S*_ is a quarnet of type 0, 1c, 1d, 1e, 1f or 2d with prune leaf *c*. This means that *c* is below a reticulation in *N*_2_ | _*S*_, and therefore, *c* has to be on side *T* or *W* of *N*_2_.

Let {*a*_*i*_, *a*_*j*_, *d*} be as before and pick *b* as an arbitrary leaf on side *B* of *N*_1_ (which exists since *N*_1_ is stack-free). Then, the quarnet *N*_1 *S*_ with *S* = {*a*_*i*_, *a*_*j*_, *b, d*}is of type 0 with the leaf *b* below the reticulation, i.e. *b* is its prune leaf. A similar argument as before then shows distinguishability, unless *b* is on side *T* or *W* of *N*_2_. We now show that this is impossible by deriving a contradiction, which will then prove Case 1. If *b* is on side *T* or *W* of *N*_2_, we can assume that the leaves on sides *B* ∪ *C* of *N*_1_ coincide with the leaves on sides *T* ∪ *W* of *N*_2_ (which also implies that |*B* | = 1 for *N*_1_). Now note that the leaves *b* and *c* are neighbours in the circular ordering of induced by *N*_1_. By our assumption, this is then also the case for *N*_2_ (where *b* and *c* are on sides *T* and *W*). Therefore, the sides *X* and *Y* (or *U* and *V*) of *N*_2_ are both empty. But then *N*_2_ is not strongly tree-child: a contradiction.

#### Case 2

*G* ≅ *H* ≅ Γ_1_. In *N*_1_, let {*a*_*i*_, *a*_*j*_} be any pair of leaves on side *A, d* any leaf on side *D*, and *σ* any leaf on side Σ ∈ {*B, C}*. Note that *a*_*i*_ and *a*_*j*_ exist since *N*_1_ is triangle-free and that *d* exists since *N*_1_ is strongly tree-child. The quarnet *N*_1 *S*_ with *S* = {*a*_*i*_, *a*_*j*_, *d, σ*} is then of type 0 or 2d, in both cases with the leaf *σ* below the reticulation (i.e. with *σ* its prune leaf). Similar reasoning (based on Corollary 2.17) as in the previous case then shows that *N*_1_ and *N*_2_ are distinguishable, unless *N*_2_ | _*S*_ is a quarnet of type 0, 1c, 1d, 1e or 1f with prune leaf *σ*. Thus, from now on we may assume that for any valid choice of *σ*, the quarnet *N*_2 *S*_ has *σ* below a reticulation, meaning that *σ* is also on sides *B* ∪ *C* of *N*_2_. Hence, *N*_1_ and *N*_2_ share the same leaves on sides *B* ∪ *C* (and therefore also on sides *A* ∪ *D*).

Now, in *N*_1_, let *a* be a leaf on side *A, b* a leaf on side *B, c* the leaf on side *C* and *d* a leaf on side *D*. Note that these leaves all exist since *N*_1_ is strongly tree-child. Furthermore, pick these four leaves such that {*a, b*} and {*c, d*} are both neighbouring pairs of leaves in the circular ordering of *X* induced by *N*_1_ (and hence also by *N*_2_). The quarnet *N*_1 *S*_ with *S* = {*a, b, c, d*} is now of type 1b, with *c* the leaf in the top middle in the planar representation of Figure 5. Recall that *N*_1_ and *N*_2_ share the same circular ordering of their leaf set *X* and that they share the same leaves on sides *B* ∪ *C* and *A* ∪ *D*. By our specific choice of the leaves in *S*, we then have that *N*_2_ | _*S*_ is either isomorphic to *N*_1_ | _*S*_, or it is a type 1b quarnet with the same circular ordering as *N*_1_ | _*S*_ but with the leaf *b* in the top middle in the planar representation of Figure 5 instead. In the latter case, *N*_1_ and *N*_2_ are distinguishable by Lemma 2.14a. Hence, we can assume that *N*_1_ | _*S*_ and *N*_2_ | _*S*_ are isomorphic, also implying that *N*_1_ and *N*_2_ share the same leaf on side *C* (and therefore also have the same leaves on side *B*).

Lastly, let *c* be the leaf on side *C* of *N*_1_ and *N*_2_. Now consider the trees-of-blobs of the networks *N*_1_ | _*S*_ and *N*_2_ | _*S*_ with *S* = *X \* {*c*}. First, assume towards a contradiction that these two trees-of-blobs are isomorphic. Then, since the two networks already share the same leaves on their side *B* and both networks have at least one leaf on their side *D*, they must share the same number of leaves on their side *D* (and therefore also on their side *A*) for the trees-of-blobs to be isomorphic. However, we already know that *N*_1_ and *N*_2_ share the same circular ordering of their leaf set *X*, showing that *N*_1_ and *N*_2_ share the same leaves on every side and that they are also ordered the same. Therefore, *N*_1_ and *N*_2_ are isomorphic: a contradiction. Thus, we may instead assume that the two trees-of-blobs are not isomorphic. But then *N*_1_ | _*S*_ and *N*_2_ | _*S*_ (and therefore also the two networks *N*_1_ and *N*_2_) are distinguishable by Corollary 2.12.

#### Case 3

*G* ≅ *H* ≅ Γ_2_. Let *t* be the leaf on side *T* of *N*_1_ and let *w* be the leaf on side *W* of *N*_1_. Note that *t* and *w* cannot be neighbours in the circular ordering of *N*_1_ and *N*_2_, otherwise *N*_1_ would not be strongly tree-child. Now consider the set *S* = X \ {*t, w*}. Then, *N*_1 *S*_ is a tree and all its induced trinets are triplet trees. If this is not the case for *N*_2_ | _*S*_, the two networks are distinguishable by Proposition 2.15, which can be used to distinguish triplet trees from trinets with a reticulation. Thus, we can assume that *N*_2 *S*_ also induces only triplet trees, i.e. *N*_2_ | _*S*_ is a tree. We now show that *t* and *w* are then also the leaves on sides *T* and *W* of *N*_2_. In particular, if neither *t* nor *w* is on sides *T* ∪ *W* of *N*_2_, we clearly have that *N*_2_ | _*S*_ is not a tree: a contradiction. On the other hand, assume w.l.o.g. that *t* is on side *T* of *N*_2_ and *w* is not on side *W* of *N*_2_. Then, since *t* and *w* are not neighbours in the circular ordering of *X* induced by *N*_2_ and since *N*_2_ is strongly tree-child, there will be some *σ*_*i*_∉ *{t, w}* on sides *X* ∪ *Y* and some *σ*_*j*_∉ *{t, w}* on sides *U* ∪ *V* of *N*_2_. Let *w*′∉ *{t, w, σ*_*i*_, *σ*_*j*_*}* then be the leaf on side *W* of *N*_2_. The trinet 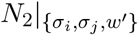 with {*σ*_*i*_, *σ*_*j*_, *w*′} ⊆ *A* will then not be a triplet tree: a contradiction. Thus, *t* and *w* are on sides *T ∪ W* of *N*_2_. Therefore, *N*_1_ and *N*_2_ share the same leaves on sides *T* and *W* (modulo swapping the two sides). Since the two networks also induce the same circular ordering of *X*, they also share the same leaves on sides *X* ∪ *Y* and *U* ∪ *V* (modulo swapping these two sets of sides). We now consider two subcases.

*Case 3.1:* |Σ| ≤ 1 *for all* Σ ∈ {*U, V, X, Y*} *for N*_1_ *and N*_2_. In this case, we have that 6, while we already showed that we may assume that |*X* | ≥ 5. When |*X* | = 6, *N*_1_ and *N*_2_ cannot be distinct while sharing the same leaves below the reticulation nodes, sharing the same leaves on sides *U* ∪ *V* and *X* ∪ *Y* (modulo swapping these two sets of sides), and sharing the same circular ordering. Hence, we get that |*X* | = 5. Then, the networks *N*_1_ and *N*_2_ are the two distinct quinnets from Figure 6 (up to relabeling the leaves). Thus, *N*_1_ and *N*_2_ are distinguishable by Lemma 2.14d.

#### Case 3.2

| Σ | ≥ 2 *for some* Σ ∈ {*U, V, X, Y*} *for N*_1_ *or N*_2_. Without loss of generality, assume that *U* 2 for *N*_1_. Now let *t* be the leaf on side *T* of *N*_1_ and set *S* = *X* \ {*t*}. Then, the tree-of-blobs of the network *N*_1 *S*_ will induce a nontrivial split that cuts off the leaves on side *U* of *N*_1_ from the rest. If the tree-of-blobs of *N*_2_ | _*S*_ is not isomorphic, the two networks are distinguishable by Corollary 2.12. Thus, we can assume they are isomorphic and the tree-of-blobs of *N*_2_|_*S*_ also induces the same split. But since *N*_1_ and *N*_2_ already share the same leaves on sides *U* ∪ *V*, this means that they share the same leaves on side *U* and on side *V*. If |*X*| ≥ 2 or *Y* ≥ 2 for *N*_1_ or *N*_2_, we can repeat this argument to obtain distinguishability between the two networks. Hence, we may assume that *X* 1 and *Y* 1 for both networks. Recall that the two networks already share the same leaves on sides *X* ∪ *Y* and that they are ordered the same. If both sets have size 1, it follows that the two networks are equal: a contradiction. Thus, for the two networks to be distinct, we assume without loss of generality that *X* = 1 and *Y* = 0 for *N*_1_, while | *X* | = 0 and |*Y* | = 1 for *N*_2_. Now, let *σ* be the leaf that is on the sides *X* ∪ *Y* of the two networks. Furthermore, let *t* be the leaf on side *T* of the networks, *w* the leaf on side *W* of the networks, *u* a leaf on side *U* of the networks, and *v* a leaf on side *V* of the networks (which both exist, since *N*_1_ is triangle-free and its side *Y* is empty, while *N*_2_ is triangle-free and its side *X* is empty). Then, *N*_1_ | _*S*_ and *N*_2_ | _*S*_ with *S* = {*σ, t, u, v, w*} are the two distinct quinnets from Figure 6 (up to relabeling the leaves). Thus, *N*_1_ and *N*_2_ are distinguishable by Lemma 2.14d.

## References

E. S. Allman, S. Petrović, J. A. Rhodes, and S. Sullivant. Identifiability of two-tree mixtures for group-based models. IEEE/ACM Transactions on Computational Biology and Bioinformatics, 8:710–722, 2010.

E. S. Allman, J. A. Rhodes, and S. Sullivant. When do phylogenetic mixture models mimic other phylogenetic models? Systematic Biology, 61:1049–1059, 2012.

E. S. Allman, J. A. Rhodes, and A. Taylor. A semialgebraic description of the general Markov model on phylogenetic trees. SIAM Journal on Discrete Mathematics, 28(2):736–755, 2014.

E. S. Allman, H. Baños, and J. A. Rhodes. NANUQ: a method for inferring species networks from gene trees under the coalescent model. Algorithms for Molecular Biology, 14:1–25, 2019.

E. S. Allman, H. Baños, and J. A. Rhodes. Identifiability of species network topologies from genomic sequences using the logDet distance. Journal of Mathematical Biology, 84(5):35, 2022.

E. S. Allman, H. Baños, J. D. Mitchell, and J. A. Rhodes. The tree of blobs of a species network: identifiability under the coalescent. Journal of Mathematical Biology, 86(1):10, 2023.

E. S. Allman, H. Baños, J. D. Mitchell, and J. A. Rhodes. TINNiK: inference of the tree of blobs of a species network under the coalescent model. Algorithms for Molecular Biology, 19(1):23, 2024a.

E. S. Allman, H. Baños, M. Garrote-López, and J. A. Rhodes. Identifiability of level-1 species networks from gene tree quartets. Bulletin of Mathematical Biology, 86(9), 2024b.

E. S. Allman, C. Ané, H. Baños, and J. A. Rhodes. Beyond level-1: identifiability of a class of galled tree-child networks. Bulletin of Mathematical Biology, 87(11):1–26, 2025a.

E. S. Allman, H. Baños, J. A. Rhodes, and K. Wicke. NANUQ+: a divide-and-conquer approach to network estimation. Algorithms for Molecular Biology, 20(1):14, 2025b.

M. Ardiyansyah. Distinguishing level-2 phylogenetic networks using phylogenetic invariants. arXiv:2104.12479, 2021.

M. L. Arnold. Natural hybridization and evolution. Oxford University Press, 1997.

H. Baños. Identifying species network features from gene tree quartets under the coalescent model. Bulletin of Mathematical Biology, 81:494–534, 2019.

E. Bapteste, L. van Iersel, A. Janke, S. Kelchner, S. Kelk, J. O. McInerney, D. A. Morrison, L. Nakhleh, M. Steel, L. Stougie, et al. Networks: expanding evolutionary thinking. Trends in Genetics, 29(8):439–441, 2013.

N. H. Barton. The role of hybridization in evolution. Molecular Ecology, 10(3):551–568, 2001.

T. Barton, E. Gross, C. Long, and J. Rusinko. Statistical learning with phylogenetic network invariants. Bulletin of the Society of Systematic Biologists, 4, 2026.

M. Casanellas, J. Fernández-Sánchez, and M. Garrote-López. SAQ: semi-algebraic quartet reconstruction. IEEE/ACM Transactions on Computational Biology and Bioinformatics, 18(6):2855–2861, 2021.

J. Cummings and B. Hollering. Computing implicitizations of multi-graded polynomial maps. Journal of Symbolic Computation, 132:102459, 2025.

A. Diop, E. L. Torrance, C. M. Stott, and L.-M. Bobay. Gene flow and introgression are pervasive forces shaping the evolution of bacterial species. Genome Biology, 23(1):239, 2022.

S. N. Evans and T. P. Speed. Invariants of some probability models used in phylogenetic inference. The Annals of Statistics, pages 355–377, 1993.

A. Francis and V. Moulton. Identifiability of tree-child phylogenetic networks under a probabilistic recombination-mutation model of evolution. Journal of Theoretical Biology, 446:160–167, 2018.

M. Frohn, N. Holtgrefe, L. van Iersel, M. Jones, and S. Kelk. Reconstructing semi-directed level-1 networks using few quarnets. Journal of Computer and System Sciences, 152:103655, 2025.

M. Frohn, N. Holtgrefe, L. van Iersel, M. Jones, and S. Kelk. Bounds on the sequence length sufficient to reconstruct binary level-1 phylogenetic networks under the CFN model. Annals of Combinatorics, 2026.

S. Ge, T. Sang, B.-R. Lu, and D.-Y. Hong. Phylogeny of rice genomes with emphasis on origins of allotetraploid species. Proceedings of the National Academy of Sciences, 96(25):14400–14405, 1999.

D. R. Grayson and M. E. Stillman. Macaulay2, a software system for research in algebraic geometry. Available at http://www2.macaulay2.com, 1992.

R. E. Green, J. Krause, A. W. Briggs, T. Maricic, U. Stenzel, M. Kircher, N. Patterson, H. Li, W. Zhai, M. H.-Y. Fritz, N. F. Hansen, E. Y. Durand, A.-S. Malaspinas, J. D. Jensen, T. Marques-Bonet, C. Alkan, K. Prüfer, M. Meyer, H. A. Burbano, J. M. Good, R. Schultz, A. Aximu-Petri, A. Butthof, B. Höber, B. Höffner, M. Siegemund, A. Weihmann, C. Nusbaum, E. S. Lander, C. Russ, N. Novod, J. Affourtit, M. Egholm, C. Verna, P. Rudan, D. Brajkovic, Željko Kucan, I. Gušic, V. B. Doronichev, L. V. Golovanova, C. Lalueza-Fox, M. de la Rasilla, J. Fortea, A. Rosas, R. W. Schmitz, P. L. F. Johnson, E. E. Eichler, D. Falush, E. Birney, J. C. Mullikin, M. Slatkin, R. Nielsen, J. Kelso, M. Lachmann, D. Reich, and S. Pääbo. A draft sequence of the Neandertal genome. Science, 328(5979):710–722, 2010.

G. Grindstaff, E. Gross, M. Hill, R. Huang, N. Riasat, and N. Vaduthala. Computing ideals of level-k Jukes-Cantor networks. forthcoming, code available at https://github.com/Rick3yHuang/Phylogenetics-Identifiability, 2025.

E. Gross and C. Long. Distinguishing phylogenetic networks. SIAM Journal on Applied Algebra and Geometry, 2(1):72–93, 2018.

E. Gross, L. van Iersel, R. Janssen, M. Jones, C. Long, and Y. Murakami. Distinguishing level-1 phylogenetic networks on the basis of data generated by Markov processes. Journal of Mathematical Biology, 83:1–24, 2021.

D. Gusfield, V. Bansal, V. Bafna, and Y. S. Song. A decomposition theory for phylogenetic networks and incompatible characters. Journal of Computational Biology, 14(10):1247–1272, 2007.

M. D. Hendy. The relationship between simple evolutionary tree models and observable sequence data. Systematic Zoology, 38(4):310–321, 1989.

M. D. Hendy and D. Penny. A framework for the quantitative study of evolutionary trees. Systematic Zoology, 38(4): 297–309, 1989.

B. Hollering and S. Sullivant. Identifiability in phylogenetics using algebraic matroids. Journal of Symbolic Computation, 104:142–158, 2021.

N. Holtgrefe, E. S. Allman, H. Baños, L. van Iersel, V. Moulton, J. A. Rhodes, and K. Wicke. Distinguishing phylogenetic level-2 networks with quartets and inter-taxon quartet distances. Bulletin of Mathematical Biology, 87(12):1–35, 2025a.

N. Holtgrefe, K. T. Huber, L. van Iersel, M. Jones, S. Martin, and V. Moulton. Squirrel: Reconstructing semi-directed phylogenetic level-1 networks from four-leaved networks or sequence alignments. Molecular Biology and Evolution, 42 (4):msaf067, 2025b.

N. Holtgrefe, K. T. Huber, L. van Iersel, M. Jones, and V. Moulton. Characterizing semi-directed phylogenetic networks and their multi-rootable variants. Theory in Biosciences, 145(1):4, 2026.

K. T. Huber, L. v. Iersel, M. Jones, V. Moulton, and L. Veenema-Nipius. When are quarnets sufficient to reconstruct semi-directed phylogenetic networks? Bulletin of Mathematical Biology, 87(10):136, 2025.

S. Huebler, R. Morris, and J. Rusinko. Constructing semi-directed level-1 phylogenetic networks from quarnets. arXiv:1910.00048, 2025.

Y. Jiao, M. An, N. Zhang, H. Zhang, C. Zheng, L. Chen, H. Li, Y. Zhang, Y. Gan, J. Zhao, H. Shang, and X. Han. Multiple third-generation recombinants formed by CRF55_01B and CRF07_BC in newly diagnosed HIV-1 infected patients in Shenzhen city, China. Virology Journal, 21(1):306, 2024.

T. H. Jukes and C. R. Cantor. Evolution of protein molecules. Mammalian Protein Metabolism, 3(24):21–132, 1969.

P. J. Keeling and J. D. Palmer. Horizontal gene transfer in eukaryotic evolution. Nature Reviews Genetics, 9(8):605–618, 2008.

M. Kimura. A simple method for estimating evolutionary rates of base substitutions through comparative studies of nucleotide sequences. Journal of Molecular Evolution, 16:111–120, 1980.

S. Kong, J. C. Pons, L. Kubatko, and K. Wicke. Classes of explicit phylogenetic networks and their biological and mathematical significance. Journal of Mathematical Biology, 84(6):47, 2022.

S. Kong, D. L. Swofford, and L. S. Kubatko. Inference of phylogenetic networks from sequence data using composite likelihood. Systematic Biology, 74(1):53–69, 2025.

D. Kosta and K. Kubjas. Geometry of symmetric group-based models. arXiv preprint arXiv:1705.09228, 2017.

J. Mallet. Hybridization as an invasion of the genome. Trends in Ecology & Evolution, 20(5):229–237, 2005.

J. Mallet. Hybrid speciation. Nature, 446(7133):279–283, 2007.

S. Martin, N. Holtgrefe, V. Moulton, and R. M. Leggett. Algebraic invariants for inferring 4-leaf semi-directed phylogenetic networks. Systematic Biology, page syaf071, 2025.

J. I. Meier, R. B. Stelkens, D. A. Joyce, S. Mwaiko, N. Phiri, U. K. Schliewen, O. M. Selz, C. E. Wagner, C. Katongo, and O. Seehausen. The coincidence of ecological opportunity with hybridization explains rapid adaptive radiation in Lake Mweru cichlid fishes. Nature Communications, 10(1):5391, 2019.

L. Nakhleh. Evolutionary phylogenetic networks: models and issues. In Problem Solving Handbook in Computational Biology and Bioinformatics, pages 125–158. Springer, 2010.

N. Patterson, D. J. Richter, S. Gnerre, E. S. Lander, and D. Reich. Genetic evidence for complex speciation of humans and chimpanzees. Nature, 441(7097):1103–1108, 2006.

J. Pekar, M. Worobey, N. Moshiri, K. Scheffler, and J. O. Wertheim. Timing the SARS-CoV-2 index case in Hubei province. Science, 372(6540):412–417, 2021.

M. Raghavan, P. Skoglund, K. E. Graf, M. Metspalu, A. Albrechtsen, I. Moltke, S. Rasmussen, T. W. Stafford, Jr, L. Orlando, E. Metspalu, M. Karmin, K. Tambets, S. Rootsi, R. Mägi, P. F. Campos, E. Balanovska, O. Balanovsky, E. Khus-nutdinova, S. Litvinov, L. P. Osipova, S. A. Fedorova, M. I. Voevoda, M. DeGiorgio, T. Sicheritz-Ponten, S. Brunak, S. Demeshchenko, T. Kivisild, R. Villems, R. Nielsen, M. Jakobsson, and E. Willerslev. Upper Palaeolithic Siberian genome reveals dual ancestry of Native Americans. Nature, 505(7481):87–91, Jan. 2014.

J. A. Rhodes, H. Baños, J. Xu, and C. Ané. Identifying circular orders for blobs in phylogenetic networks. Advances in Applied Mathematics, 163:102804, 2025.

L. H. Rieseberg. Hybrid origins of plant species. Annual review of Ecology and Systematics, 28(1):359–389, 1997.

E. B. Sessa, E. A. Zimmer, and T. J. Givnish. Unraveling reticulate evolution in North American Dryopteris (dryopteridaceae). BMC Evolutionary Biology, 12:1–24, 2012.

C. Solís-Lemus and C. Ané. Inferring phylogenetic networks with maximum pseudolikelihood under incomplete lineage sorting. PLoS Genetics, 12(3):e1005896, 2016.

S. M. Soucy, J. Huang, and J. P. Gogarten. Horizontal gene transfer: building the web of life. Nature Reviews Genetics, 16 (8):472–482, 2015.

B. Sturmfels and S. Sullivant. Toric ideals of phylogenetic invariants. Journal of Computational Biology, 12(4):457–481, 2005.

S. Sullivant. Algebraic statistics, volume 194. American Mathematical Society, 2023.

S. Sullivant. Phylogenetic network models as directed acyclic graphs. arXiv:2507.23056, 2025.

G. J. Szöllősi, A. A. Davín, E. Tannier, V. Daubin, and B. Boussau. Genome-scale phylogenetic analysis finds extensive gene transfer among fungi. Philosophical Transactions of the Royal Society B: Biological Sciences, 370(1678):20140335, 2015.

S. A. Taylor and E. L. Larson. Insights from genomes into the evolutionary importance and prevalence of hybridization in nature. Nature Ecology & Evolution, 3(2):170–177, 2019.

X.-X. Wang, C.-H. Huang, D. F. Morales-Briones, X.-Y. Wang, Y. Hu, N. Zhang, P.-G. Zhao, X.-M. Wei, K.-H. Wei, X. Hemu, N.-H. Tan, Q.-F. Wang, and L.-Y. Chen. Phylotranscriptomics reveals the phylogeny of Asparagales and the evolution of allium flavor biosynthesis. Nature Communications, 15(1):9663, 2024.

S. J. Westenberger, C. Barnabé, D. A. Campbell, and N. R. Sturm. Two hybridization events define the population structure of Trypanosoma cruzi. Genetics, 171(2):527–543, 2005.

M. Worobey, M. Gemmel, D. E. Teuwen, T. Haselkorn, K. Kunstman, M. Bunce, J.-J. Muyembe, J.-M. M. Kabongo, R. M. Kalengayi, E. Van Marck, M. T. P. Gilber, and S. M. Wolinsky. Direct evidence of extensive diversity of HIV-1 in Kinshasa by 1960. Nature, 455(7213):661–664, 2008.

J. Xu and C. Ané. Identifiability of local and global features of phylogenetic networks from average distances. Journal of Mathematical Biology, 86(1):12, 2023.

Y. Yu, C. Than, J. H. Degnan, and L. Nakhleh. Coalescent histories on phylogenetic networks and detection of hybridization despite incomplete lineage sorting. Systematic Biology, 60(2):138–149, 2011.

